# Cross-attractor modeling of resting-state functional connectivity in psychiatric disorders

**DOI:** 10.1101/2022.10.29.514373

**Authors:** Yinming Sun, Mengsen Zhang, Manish Saggar

## Abstract

Resting-state functional connectivity (RSFC) is altered across various psychiatric disorders. Brain network modeling (BNM) has the potential to reveal the neurobiological underpinnings of such abnormalities by dynamically modeling the structure-function relationship and examining biologically relevant parameters after fitting the models with real data. Although innovative BNM approaches have been developed, two main issues need to be further addressed. First, previous BNM approaches are primarily limited to simulating noise-driven dynamics near a chosen attractor (or a stable brain state). An alternative approach is to examine multi(or cross)-attractor dynamics, which can be used to better capture non-stationarity and switching between states in the resting brain. Second, previous BNM work is limited to characterizing one disorder at a time. Given the large degree of co-morbidity across psychiatric disorders, comparing BNMs across disorders might provide a novel avenue to generate insights regarding the dynamical features that are common across (vs. specific to) disorders. Here, we address these issues by (1) examining the layout of the attractor repertoire over the entire multi-attractor landscape using a recently developed cross-attractor BNM approach; and (2) characterizing and comparing multiple disorders (schizophrenia, bipolar, and ADHD) with healthy controls using an openly available and moderately large multimodal dataset from the UCLA Consortium for Neuropsychiatric Phenomics. Both global and local differences were observed across disorders. Specifically, the global coupling between regions was significantly decreased in schizophrenia patients relative to healthy controls. At the same time, the ratio between local excitation and inhibition was significantly higher in the schizophrenia group than the ADHD group. In line with these results, the schizophrenia group had the lowest switching costs (energy gaps) across groups for several networks including the default mode network. Paired comparison also showed that schizophrenia patients had significantly lower energy gaps than healthy controls for the somatomotor and visual networks. Overall, this study provides preliminary evidence supporting transdiagnostic multi-attractor BNM approaches to better understand psychiatric disorders’ pathophysiology.

## Introduction

Resting-state functional connectivity (RSFC) is observed to be altered across various psychiatric disorders, including schizophrenia, bipolar disorder, and attention deficit hyperactivity disorder (ADHD) (Baker et al., 2019; Friston et al., 2016; Khadka et al., 2013; Konrad and Eickhoff, 2010; McCarthy et al., 2013; Perry et al., 2019; Xia et al., 2018). Likewise, structural connectivity (SC) based on diffusion-weighted imaging (DWI) has also revealed significant deviations across patient populations (Favre et al., 2019; Friston et al., 2016; Kelly et al., 2018; Konrad and Eickhoff, 2010; van Ewijk et al., 2012). Here, we argue for using a brain network modeling (BNM) approach that captures structure-function relationships to better characterize disorder-specific findings across modalities. BNM models are nonlinearly dependent on structure, which allows them to capture additional variances in function through the synergistic effect of the structural connectome and biophysical parameters (Breakspear, 2017; Deco et al., 2011).

A vital benefit of the BNM approach is that it allows for examining differences in modeled physiological parameters (e.g., inhibitory synaptic strength) and generates concrete hypotheses regarding the neurobiological differences associated with psychiatric disorders. Along this line of thinking, previous studies have shown that RSFC can be partially predicted using SC directly or via modeling approaches (Hagmann et al., 2008; Honey et al., 2009; Kringelbach and Deco, 2020; Schirner et al., 2018). In terms of modeling psychiatric disorders, one BNM study with schizophrenia patients demonstrated how an increase in the regional excitation/inhibition (E/I) ratio led to an increase in functional connectivity, especially in the frontal-parietal network (Yang et al., 2016). Another study on autism patients showed that increased recurrent E/I explained abnormalities in both somatosensory regions and association cortices (Park et al., 2021). Finally, a study on ADHD patients found abnormalities in a model parameter linked to elevated regional oscillations and identified two subgroups of patients differing in personality traits (Iravani et al., 2021).

While previous modeling studies have advanced our understanding of psychiatric disorders, several key issues can be better addressed. First, previous applications of BNM in clinical populations were primarily limited to simulating noise-driven (or stochastic) dynamics near a chosen attractor (or a stable brain state (Gustavo Deco et al., 2013; Demirtaş et al., 2019); see (Cabral et al., 2017) for more details). As an alternative approach, multi(or cross)-attractor examinations can be used to better capture non-stationarity and switching between states in the resting brain (Deco and Jirsa, 2012; Freyer et al., 2012; Hansen et al., 2015; Zhang et al., 2023, 2022). Second, previous BNM work was primarily limited to characterizing one disorder at a time. Given the large degree of co-morbidity across psychiatric disorders and the recent push in the field toward examining biological features across disorders, comparing BNMs across multiple disorders might provide a novel avenue to generate insights regarding the common dynamical features across disorders vs. specific to each disorder.

Here, we address both of these issues by (1) examining the layout of the attractor repertoire over the entire multi-attractor landscape using our recently developed cross-attractor BNM approach (Zhang et al., 2022); and (2) characterizing and comparing multiple disorders (schizophrenia, bipolar, and ADHD) with healthy controls using an openly available and moderately large multimodal dataset (LA5c) from the UCLA Consortium for Neuropsychiatric Phenomics (Poldrack et al., 2016).

Using data from undiagnosed adults from the Human Connectome Project (HCP), we have recently shown that real data derived RSFC can be accurately explained by the set of possible transitions between all attractors, termed cross-attractor coordination matrix (Zhang et al., 2022). In contrast to single-attractor based approaches for modeling the RSFC (G. Deco et al., 2013; Demirtaş et al., 2019; Ghosh et al., 2008), our modeling approach quantifies how well brain regions co-fluctuate across all possible attractor states. This deterministic approach provides a summarizing metric of the attractors landscape and has been shown to be especially effective at explaining functional connections across hemispheres seen in the real data (Zhang et al., 2022).

Moreover, we also defined the concept of the “energy gap” between attractor states for characterizing the potential costs of state switching. Here, to capture individual differences in neurobiology across disorders, we varied model parameters for local excitation and inhibition and the global coupling between brain regions. The optimal individual combination of the parameters was determined based on how well the cross-attractor coordination matrix fits with the experimentally measured RSFC. Based on the optimal model configuration, we calculated the associated global energy gap measures for each subject following our previous study (Zhang et al., 2022). Since the abnormalities may localize to specific brain regions or functions (Baker et al., 2019; Ishida et al., 2023; Kebets et al., 2019), we also examined energy gap metrics averaged across regions of canonical resting-state networks (Yeo et al., 2011). The distribution of each model parameter and the global and network-specific energy gap measures were compared across the groups to identify disorder-specific abnormalities.

We expect the model fitness to be similar across participant groups. Based on previous findings of inhibitory neuron deficits, we hypothesize that parameters for local inhibition would be affected for schizophrenia and bipolar patients (Benes and Berretta, 2001; Lewis et al., 2012). We also expect higher values for energy gap measures in schizophrenia patients since they are associated with cognitive deficits and more severe psychopathology, which may result in more difficult transitions between attractor states. Regarding network-specific energy gap effects, we hypothesize the default mode network to show significant abnormalities given the large amount of evidence for its role in various psychiatric disorders (Baker et al., 2019; Bluhm et al., 2007; Whitfield-Gabrieli and Ford, 2012). Since ADHD has been associated with deficits in attention networks (McCarthy et al., 2013), we expect more significant abnormalities for energy gap metrics in the dorsal and ventral attention networks.

Overall, we aim to better characterize the commonality and differences between psychiatric disorders using a multi(cross)-attractor BNM model and a transdiagnostic dataset.

## Methods

### Participants

The LA5c dataset was available through the OpenNeuro website, with further details presented elsewhere (Poldrack et al., 2016). In brief, adults ages 21 to 50 years were recruited from the Los Angeles area as part of the Consortium for Neuropsychiatric Phenomics. All participants gave informed consent and were either healthy (HLTY) or had a clinical diagnosis of schizophrenia (SCHZ), bipolar disorder (BPLR), or attention deficit hyperactivity disorder (ADHD). The downloaded dataset included 130 HLTY, 50 SCHZ, 49 BPLR, and 43 ADHD.

### Neuroimaging data description and analysis

The MRI data were collected using two 3T Siemens Trio scanners and included a 1 mm T1 scan with MPRAGE sequence, a 2 mm 64-direction DWI scan with one shell (b = 1000 s/mm^2^), and a 4 mm echo planar imaging (EPI) resting state fMRI (rsfMRI) scan with 2 s TRs and lasting 304 seconds.

rsfMRI was preprocessed using the automated workflow from fMRIprep, which is elaborated elsewhere (Esteban et al., 2019). The output fMRI in standard ‘MNI152NLin6Asym’ space was further processed by removing (censuring) volumes with a framewise displacement of 0.5 mm or higher, and regressing out six motion parameters (three translational and three rotational) from the remaining good frames. Any subject with less than 80% good frames was excluded from further analysis. To generate the RSFC, we used the Desikan-Killiany (DK) atlas (Gustavo Deco et al., 2013; Zhang et al., 2022) with 66 parcels (excluding left and right insula). The parcellation matches our previous study on HCP subjects and has been shown to be a good compromise between biological realism and computation runtime (Gustavo Deco et al., 2013; Zhang et al., 2022). For easier comparison of connectivity between corresponding regions of the two hemispheres, the sorting order of the parcels in the two hemispheres was flipped (sorting order shown in Supplemental Fig. 10b). This sorting order often results in a top right to bottom left diagonal in the connectivity matrix due to strong connectivity between matching regions of the two hemispheres (Gustavo Deco et al., 2013; Zhang et al., 2022). Whole brain rsfMRI time series were first averaged for voxels part of each parcel defined based on the subject DK atlas in standard ‘MNI152NLin6Asym’ space created via Freesurfer as part of the fMRIprep workflow. The parcel level time series were then correlated between each pair of parcels to generate the RSFC matrix.

For the analysis of diffusion scans, MRtrix3 was used for preprocessing, computing fixel-based values, and generating the probabilistic streamlines (Tournier et al., 2019). Preprocessing included removing random noise and ringing artifacts, removing distortion caused by eddy currents using the eddy functionality from FSL (Andersson et al., 2003), and applying bias field correction using the N4 algorithm of ANTs (Gustavo Deco et al., 2013; Wong and Wang, 2006). Subsequently, tissue response functions representing single-fiber white matter, gray matter, and cerebral spinal fluid (CSF) were computed and used for estimating the fiber orientation distribution (FOD) based on the multi-tissue constrained spherical deconvolution approach (Tournier et al., 2007). To generate an anatomically constrained tractography (ACT) (Gustavo Deco et al., 2013), the structural T1 was co-registered to the DWI scan and was used to generate tissue segmentation of cortical gray matter, subcortical gray matter, white matter, CSF, and pathological tissue. Using the resulting tissue segmentations, 10 million probabilistic streamlines were generated with the MRtrix3 default iFOD2 algorithm by seeding at the gray and white matter interface. To remove biases in the whole-brain tract generation process, the number of streamlines was down-sampled to 1 million based on the SIFT algorithm (Smith et al., 2013). The co-registered subject-specific DK atlas was used to generate the SC by counting the number of streamlines between each parcel pair and normalizing by the average of the two parcel volumes. The resulting raw SC was further processed by setting the diagonal elements to zero and then normalizing by the value of the total connectivity from the parcel with the highest total connectivity (i.e., infinity normalized, Equation 4). For quality control based on outlier detection, the similarity between the SC of all subjects was quantified with Spearman’s correlation between the lower diagonal entries of the symmetrical matrix. Any subject with a similarity score of less than three standard deviations from the group mean was excluded from further analyses.

After excluding subjects with poor fMRI and outlier DWI connectivity, there were 104 HLTY, 29 SCHZ, 41 BPLR, and 31 ADHD participants. The demographics of the participants and their average framewise displacement (FD), are shown in Table 1. Significant group differences were observed for age, site (scanner), and average FD. Post-hoc, no significant paired comparison difference was observed for the overall age effect (ANOVA F = 3.04, p = 0.03). For the overall site effect (Pearson’s 𝒳^2^ = 22.7, p < 0.001), the HLTY group had a larger percentage of participants from site 1 and a lower percentage from site 2 than expected (|standardized residuals| = 4.7). For the effect of average FD (ANOVA F = 5.21, p = 0.002), the HLTY group had a significantly lower value than both the SCHZ group (p = 0.008) and the BPLR group (p = 0.028).

**Table 1:** Participant demographics.

| Group | N | Age* | Sex (Male/Female) | Site* (1/2) | Average Framewise Displacement (FD) * |
| --- | --- | --- | --- | --- | --- |
| Healthy (HLTY) | 104 | 30.8±8.4 | 53/51 | 83/21 | 0.12±0.05 |
| Schizophrenia (SCHZ) | 29 | 34.7±9.5 | 22/7 | 15/14 | 0.16±0.05 |
| Bipolar Disorder (BPLR) | 41 | 35.0±8.4 | 22/19 | 21/20 | 0.15±0.07 |
| Attention Deficit Hyperactivity Disorder (ADHD) | 31 | 32.3±8.4 | 18/13 | 13/18 | 0.13±0.06 |

\* Significant group difference based on omnibus test

### Overview of the brain network model

The implementation details of our BNM model tailored for generating cross-attractor-based RSFC (Figure 1) have been presented elsewhere (Zhang et al., 2022). Briefly, the model has parameters that can be linked with biological quantities (i.e., an ensemble of leaky integrate- and-fire neurons receiving uncorrelated noisy inputs) as in the Wong-Wang model (Wong and Wang, 2006) and is modified such that the transfer function from input current to firing rate is similar to the Wilson-Cowen model (Wilson and Cowan, 1972), which captures a more diverse range of local dynamics.

**Figure 1.**
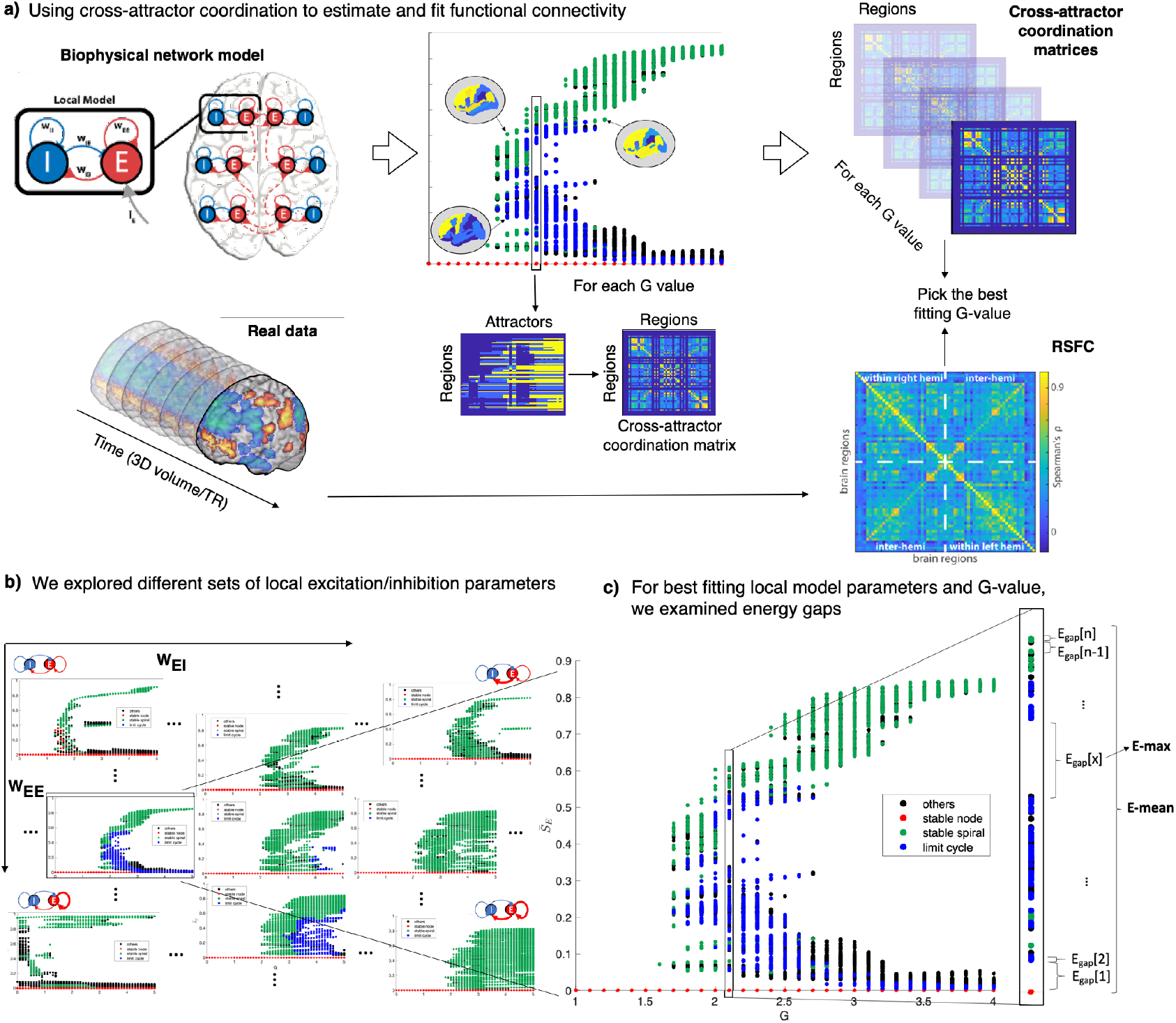
Overview of cross-attractor-based model: Panel a) shows how our model is used to calculate the cross-attractor coordination matrix from the set of all attractors, which is then fitted with the real data derived RSFC. Panel b) shows a set of bifurcation plots generated from 88 different local configurations. Panel c) shows a zoomed-in version of one bifurcation plot to illustrate what the maximum and mean energy gap metrics (E-max, E-mean) represent.

Equations 1 and 2 describe how the synaptic activity or fraction of open synaptic channels of excitatory 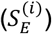 and inhibitory 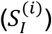 populations evolve in brain region *i*. For each region 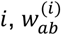 represent the local coupling from population *a* to *b, H*_*p*_ is the sigmoidal transfer function for population 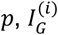 represent the global external input to the region *i* and 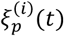 is the intrinsic noise for population *p*. The decay time constants (*τ*), kinetic parameters (*γ*), are fixed for each population across regions, while the noise scaling constant (*σ*) is constant across both populations (see Table 2).

**Table 2:** Fixed model parameter values. For details on the choice of values, please refer to (Zhang et al., 2022) and previous literature (Gustavo Deco et al., 2013; Wong and Wang, 2006).

| Parameter | Value | Interpretation |
| --- | --- | --- |
| $\tau_E$ | 0.1 (s) | Decay time of NMDA receptor |
| $\tau_I$ | 0.01 (s) | Decay time of GABA receptor |
| $\gamma_E$ | 0.641 | Kinetic parameter of excitatory population |
| $\gamma_I$ | 1 | Kinetic parameter of inhibitory population |
| $\sigma$ | 0.01 | Amplitude of intrinsic noise |
| $a_E$ | 310 (nC<sup>-1</sup>) | Slope of the near-linear segment of sigmoidal curve (excitatory population) |
| $b_E$ | 125 Hz | Middle of near-linear segment of sigmoidal curve (excitatory population) |
| $d_E$ | 0.16 (s) | Smoothness of the corner bends of sigmoidal curve (excitatory population) |
| $a_I$ | 615 (nC<sup>-1</sup>) | Slope of the near-linear segment of sigmoidal curve (inhibitory population) |
| $b_I$ | 177 (Hz) | Middle of near-linear segment of sigmoidal curve (inhibitory population) |
| $d_I$ | 0.087 (s) | Smoothness of the corner bends of sigmoidal curve (inhibitory population) |
| $r_{max}$ | 500 (Hz) | Maximal firing rate |
| $w_{II}$ | 0.05 (nA) | Inhibitory-to-inhibitory coupling |
| $I_I$ | 0.1 (nA) | Global input current to inhibitory population |

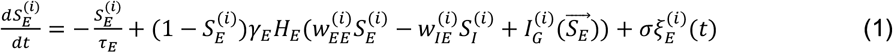

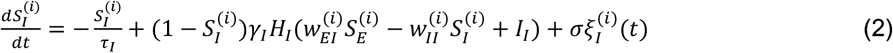

Equation 3 describes how the global input current 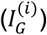 to the excitatory population of each region *i* is affected by the activity of all other regions *j*, which is scaled by the structural connectivity (*C*_*ij*_) and global coupling variable *G*. To generate *C*_*ij*_, diagonal elements of the raw SC matrix were first set to 0. Then, the matrix elements were normalized by the maximum of the column sums (Equation 4).

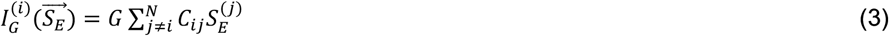

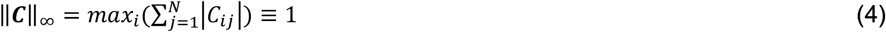

Equation 5 describes the sigmoidal transfer function that converts input current *x* for a neuronal population (*p*) into output firing rate *H*_*p*_(*x*), with constants *a, b, d*, and *r*_*max*_ that defines the shape of the sigmoidal curve.

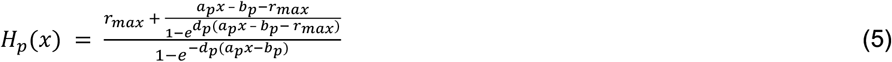

### Computing attractor states

Due to the limited quality of the diffusion scans and previous studies showing the effectiveness of using a group average SC for the BNM modeling (Iravani et al., 2021; Zhang et al., 2022), we decided to use the group average SC for this study. Using a group-specific SC instead of one for all groups allowed for preserving group variations in SC in the subsequent analysis. The DK parcellation was chosen for comparison with previous works (Gustavo Deco et al., 2013; Kringelbach and Deco, 2020; Zhang et al., 2022) and its optimal trade-off in terms of regional homogeneity and computational efficiency. To encompass the possible optimal fits across groups and individuals within each group, a sizeable regular grid of 88 local configurations of (w_EE_, w_EI_), spanning from 0.5 to 4 for w_EE_ with a step size of 0.5 (8 combinations) and spanning from 0.5 to 3 with a step size of 0.25 for w_EI_ (11 combinations), were each used for computing cross-attractor coordination. Further, the global coupling parameter *G* was varied from 0 to 5 with 0.1 increments. Similar to the bounds set for w_EE_ and w_EI_, the *G* range was set to capture an extensive range of values for fitting individual differences in RSFC. In contrast, the increment size was set to minimize the number of run configurations but still be able to track the rate of change in the dynamic landscape. For each specific configuration of global coupling (*G*), connectivity matrix (*C*), local excitation (*w*_*EE*_), and local inhibition (*w*_*EI*_), the set of fixed points was determined by recursively searching for steady-state solutions or zeroes of equations 1 and 2 from an initial set of guesses and then making new guesses based on the solutions found. The recursive process was repeated until a preset number of zeroes were found, or a specific recursion depth limit was reached. Further details of the algorithm are presented in our previous work (Zhang et al., 2022). To determine the set of stable fixed points or attractors from all the fixed points, we first classified the fixed points based on the Jacobian of the solution. We then performed an additional perturbation test around each fixed point for verification. For this verification step, the integration step was set at 0.001 second. Numeric integration is not used for any other part of our analysis. Numeric integration is not used for any other part of our analysis. Figures 1b and 1c show example bifurcation diagrams corresponding to the different local configurations, with all the attractors labeled by type: stable nodes, stable spirals, and limit cycles. The remaining fixed points, labeled as others, are unstable and not attractors. Since the group average SC was used in this study, 88 bifurcation diagrams (one for each local configuration), portraying all possible attractor states, were generated for each group.

### Computing cross-attractor coordination

Computation of the cross-attractor coordination is identical to that of our previous work (Zhang et al., 2022), described in Figure 2 and summarized as follows. For each specific configuration, an attractors-by-regions matrix, or an attractor repertoire *A*(*G, C, w*_*EE*_, *w*_*EI*_), was assembled by combining the values of each attractor 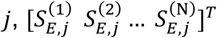 as a column vector, where the element 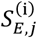 is the value of region *i* for attractor *j* (Figure 2a). The regional 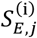 values across all matrix elements of *A* were then used to generate a distribution, from which bins were defined with the local minima and two edges as boundaries (see Figure 2c). A discretized attractor repertoire *Â* was created by assigning the original matrix values with the bin number they were part of. The discretized attractor values for each region (Figure 2d) were subsequently correlated with each other, resulting in a symmetrical regions-by-regions matrix *P*, termed the cross-attractor coordination matrix (Figure 2f). An intuitive understanding of the cross-attractor coordination matrix is that it quantifies the level of coordinated activity between regions across all the attractors. A higher matrix value between two regions means they are more coordinated in their activity changes when switching between attractor states.

**Figure 2.**
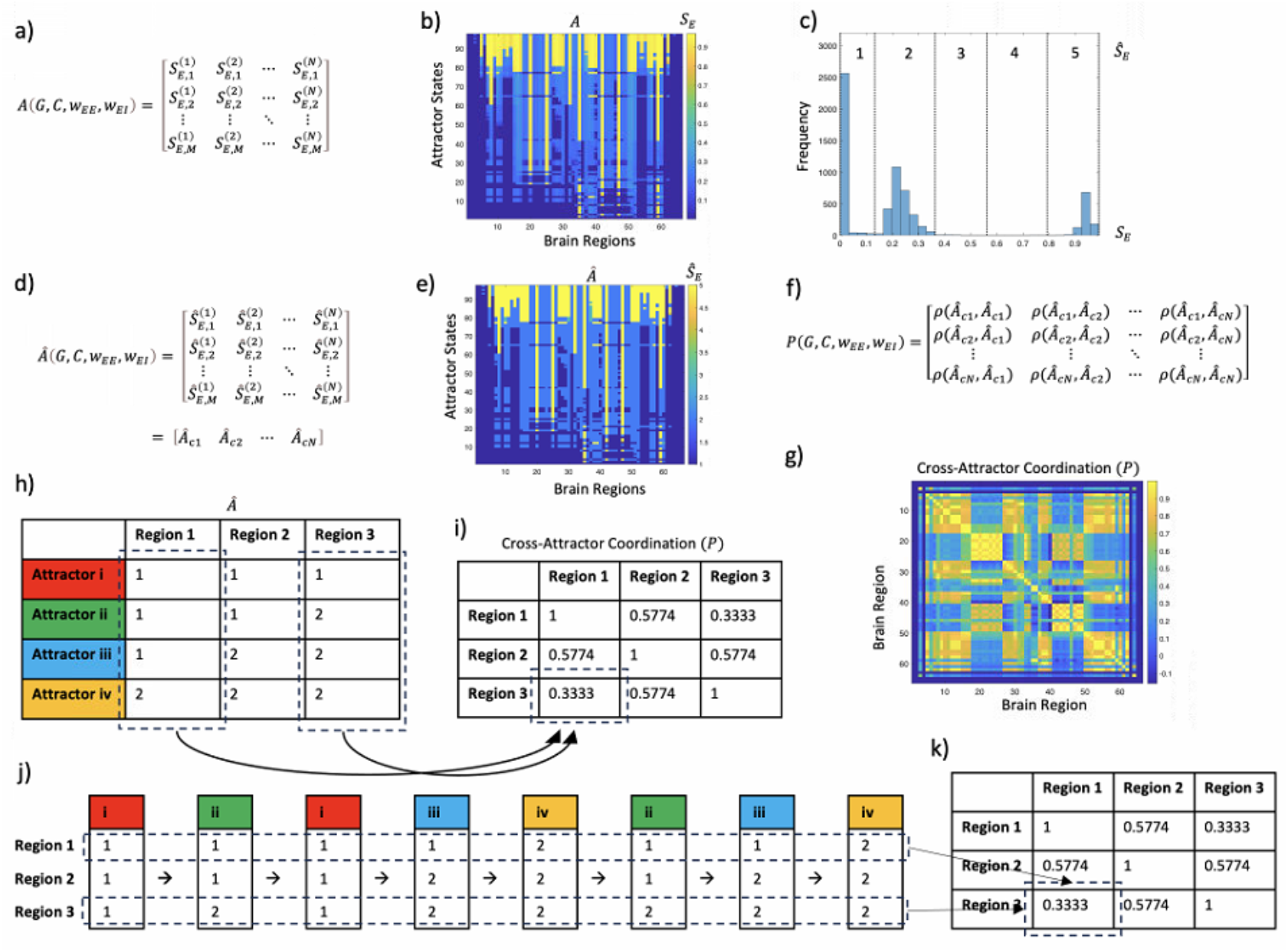
Cross-attractor coordination matrix and its potential relation to functional connectivity. Cross-attractor coordination matrix for a specific configuration of global coupling (G), connectivity matrix (C), local excitation (w_EE_), and local inhibition (w_EI_) is computed from the list of M attractors and N regions (i.e., M by N matrix A(G, C, w_EE_, w_EI_)) by first discretizing the attractor values through binning of the distribution of S_E_ across all attractors and brain regions, resulting in a discretized M by N matrix A,(G, C, w_EE_, w_EI_), and then computing the Spearman’s correlation between the columns of Â, which are the attractor state values for each brain region, to produce a N by N cross-attractor coordination matrix P(G, C, w_EE_, w_EI_). Further explanations of each step are provided in the text. The general matrix form of the three matrices A, Â, and P are shown in subpanels a), d), and f) respectively. 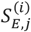 refers to the S_E_ value for region i and attractor j; 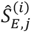 refers to the discretized Ŝ_E_ value for region i and attractor j; ρ(Â_ci_, Â_cj_) refers to the Spearman correlation between columns i and j of Â. A real data example of the three matrices are shown in subpanels b), e), and g) respectively. Subpanel c) illustrates the discretization process for an exemplar data. Subpanel h) shows a simplified toy example (4 attractors, 3 regions, and 2 discretized states) illustrating how A, is used to calculate P. With the toy example, subpanels j) and k) illustrate how the cross-attractor matrix is conceptually related to the traditional definition of functional connectivity (i.e., the correlation between the neural time series of brain regions). Specifically, assuming the brain cycles through the attractor states with an equal probability of traversing each attractor, then as time increases, the functional connectivity k) becomes the cross-attractor coordination. For simplicity of illustration, the example time series was deliberately chosen with an equal occurrence of each attractor (i.e., 2 times), which would be true of any sequence satisfying the assumptions as the time approaches infinity.

The cross-attractor coordination matrix can be correlated with the real RSFC because functionally connected regions are expected to have co-fluctuating attractor states (i.e., similar values for the same attractor). Moreover, the cross-attractor coordination matrix may be related to the classical definition of functional connectivity (i.e., the correlation between the neural time series of brain regions) if the attractor states can be seen as states that the brain traverses through during the resting state. In fact, if the probability of traversing each attractor is the same, then as time increases, the functional connectivity would approach the cross-attractor coordination matrix (see Figure 2 j,k). It is important to note that these attractor states are not the same as dynamical functional network states, as shown in previous literature (Zhang et al., 2023).

### Individual RSFC fitting with cross-attractor coordination

The optimal configuration in terms of <*G, w*_*EE*_, and *w*_*EI*_> for each participant, was determined by comparing the Spearman’s correlation between the real RSFC and the cross-attractor coordination matrix associated with each configuration (Zhang et al., 2022) (Figure 1a). This approach to evaluating fitness is commonly used in the literature (G. Deco et al., 2013; Demirtaş et al., 2019; Park et al., 2021; Wang et al., 2019). Since 88 different local configurations (8 *w*_*EE*_ x 11 *w*_*EI*_) were modeled with a *G* spanning between 1 and 5 and a step size of 0.1 (51 possibilities), the best fitting cross-attractor coordination matrix (i.e., highest Spearman’s correlation) was found by comparing 4488 model combinations. To prevent the fitting of RSFC to a cross-attractor coordination matrix associated with unrealistic attractor repertories with large gaps between separate sub-repertoires, the maximum energy gap allowed for fitting was set to be no greater than 0.2 based on evidence from our previous work (Zhang et al., 2022). Specifically, the max energy gap for healthy subjects fitted to the model without an energy constraint was mainly less than 0.2, and the loss in model fitness with the constraint only becomes evident above that threshold.

### Energy gap calculations

Energy gap measures were calculated at the optimal (*G, w*_*EE*_, and *w*_*EI*_) for each subject. Based on our previous study (Zhang et al., 2022), the energy gap was defined by calculating the average S_E_ across all regions for each attractor and then taking the difference between adjacent attractors after sorting by the global average S_E_ values. While other ways of describing the multi-attractor landscape are possible, this simple summary metric is an intuitive way of relating the attractors based on the difference in the fraction of open post-synaptic ion channels (i.e., S_E_). A larger energy gap means that more energy is required to open the additional ion channels in the higher attractor state relative to the lower attractor state. The maximum and mean energy gap (E-max, E-mean) were then calculated from the distribution of energy gaps for each subject (Figure 1c). Since these measures were based on the global average S_E_, we referred to them as global energy gap measures.

Additionally, we also examined network-specific energy gaps. Network-specific energy gaps between adjacent attractors were defined as the difference between the S_E_ averaged across brain regions that are part of specific canonical networks (Yeo et al., 2011). Each parcellated region of the DK atlas was assigned to a particular network based on which network had the most vertices in the region and included more than 25% of the vertices (Supplemental Fig. 10). The attractors were sorted based on the global S_E_ since attractor states themselves were defined based on the global average.

### Group comparison of individual values

The optimal *G*, the fitness correlation (Spearman’s ρ), *w*_*EE*_, *w*_*EI*_, *w*_*EE*_/*w*_*EI*_ ratio, E-max, and E-mean were compared across the clinical groups with the Kruskal-Wallis test after regressing out covariates for age, sex, site (scanner), and average FD. Post-hoc rank-sum tests with Tukey correction were used to determine significant paired differences if there was an overall group effect. Non-parametric rank-based tests were used since the distributions of the variables were not normal, and the tests are more robust to uneven distributions of individual values.

## Results

### Description of attractor landscapes

The bifurcation diagram of S_E_ with changing values of *G* agrees with previous studies (Gustavo Deco et al., 2013; Zhang et al., 2022). Key characteristics include an initial jump from a single stable state to multiple stable states or multistability, and a second bifurcation that splits the set of attractors to a lower and higher arm (Figure 1). The arms then gradually collapse toward two extreme states with further increases in *G*.

Bifurcation diagrams for the healthy group are shown in Supplemental Fig. 15 to 22. To better illustrate how the landscape changes in attractor type and numbers, Supplemental Fig. 23 to 30 show the total number of attractors and the number of attractors of each type for the healthy group. As *w*_*EE*_ increases, the gap between the first and second major bifurcation value of *G* is widened. The attractors also span a wider range of S_E_ values for higher *G* values. As w_EI_ increases, the entire bifurcation diagram shifts toward a higher *G*, including the *G* value at the first and second major bifurcation. The total number of possible attractors across all *G* increases while the number of limit cycle solutions decreases and occurs at higher *G* values. The same general trends hold for the bifurcation plots of all patient groups.

### Individual model fitting is similar across groups

Figure 3a shows the distribution of correlation (Spearman’s ρ) between the cross-attractor coordination matrix and real RSFC for each participant group. The individual fitting of model parameters resulted in very similar correlation values for each of the 4 groups, which were 0.44 (0.09) for HLTY, 0.48 (0.08) for SCHZ, 0.47 (0.12) for BPLR, and 0.42 (0.09) for ADHD. Overall, model fitting was at par with previous studies, which had fitness correlation values between 0.4 and 0.5 (Gustavo Deco et al., 2013; Iravani et al., 2021; Park et al., 2021; Wang et al., 2019). After regressing out the effect of age, sex, site (scanner), and average FD, there was no significant group effect for the fitness correlation based on a 1-way Kruskal-Wallis test (𝒳^2^ = 3.29, p = 0.4), suggesting that individual fitness was similar across both healthy controls and patients.

**Figure 3.**
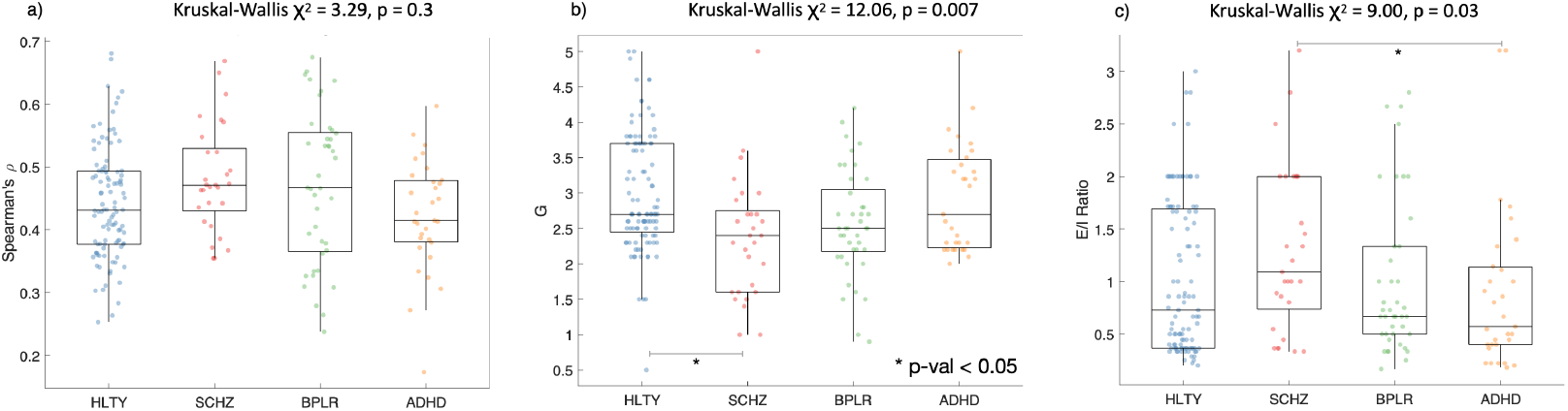
Model parameter fitting: Boxplots showing the distribution across group participants for a) model fitness (Spearman’s ρ), b) global coupling (G) value, and c) excitation to inhibition ratio (w_EE_/w_EI_). The subpanel titles show the Kruskal-Wallis test results for group comparisons after controlling for age, sex, site, and average FD. For comparisons with a significant overall group effect, post hoc ranksum tests were done between each group pair. Significant paired differences after Tukey correction (* p-val < 0.05) are marked by a line and star.

Example fitting results for selected participants are shown in Supplemental Fig. 1 to 4. Stability of fitness values (Spearman’s rho) was observed around the maximum, suggesting reliable fitting. The distribution of all model parameters for each group is shown in Supplemental Fig. 5 to 8. The correlation with RSFC was higher for cross-attractor coordination than for group SC for nearly all individuals (see Supplemental Fig. 9). As expected, the cross-attractor coordination matrix of individuals is correlated to the group SC, which was used as part of the BNM model.

### Global coupling is decreased in schizophrenia

The distribution of optimal *G* associated with the correlation values is shown in Figure 3b. The average *G* value was 3.0 (0.8) for HLTY, 2.4 (0.9) for SCHZ, 2.6 (0.8) for BPLR, and 2.9 (0.8) for ADHD. These values are close to where the fitted bifurcation diagram starts to split into a high and low sub-repertoire of attractors, or what Deco and colleagues refer to as the edge of the second bifurcation or criticality (Deco et al., 2013). It is also where the total number of attractors is typically the largest (see Supplemental Fig. 23 to 30). After regressing out the effect of age, sex, site (scanner), and average framewise displacement, there was a significant group effect for the optimal *G* based on a 1-way Kruskal-Wallis test (𝒳^2^ = 12.06, p = 0.007). Post-hoc paired comparisons show that *G* was significantly higher for HLTY than SCHZ after correction for multiple comparisons.

### Optimal local configuration differs across groups

There is wide individual variability in terms of local configuration (w_EE_, w_EI_) within each group. For the HLTY group, the average w_EE_ is 1.83 (1.04), the average w_EI_ is 2.10 (0.63), and the average w_EE_/w_EI_ ratio is 1.02 (0.72). For the SCHZ group, the average w_EE_ is 1.88 (1.06), the average w_EI_ is 1.76 (0.85), and the average w_EE_/w_EI_ ratio is 1.30 (0.78). For the BPLR group, the average w_EE_ is 1.61 (0.86), the average w_EI_ is 2.04 (0.83), and the average w_EE_/w_EI_ ratio is 1.00 (0.74). For the ADHD group, the average w_EE_ is 1.71 (1.09), the average w_EI_ is 2.20 (0.51), and the average w_EE_/w_EI_ ratio is 0.89 (0.77). After regressing out the effect of age, sex, site (scanner), and average framewise displacement, there was a significant group effect for the w_EE_/w_EI_ ratio. Post-hoc comparisons show that the ADHD group has a lower w_EE_/w_EI_ ratio than the SCHZ group.

### The global energy gap is similar across groups

The distribution of the global E-max is shown in Figure 4a. The group average E-max was 0.09 (0.04) for HLTY, 0.09 (0.03) for SCHZ, 0.08 (0.03) for BPLR, and 0.09 (0.04) for ADHD. The distribution of global E-mean is shown in Supplemental Fig. 11a. The group average E-mean was 0.02 (0.01) for HLTY, 0.02 (0.01) for SCHZ, 0.02 (0.01) for BPLR, and 0.02 (0.01) for ADHD.

**Figure 4.**
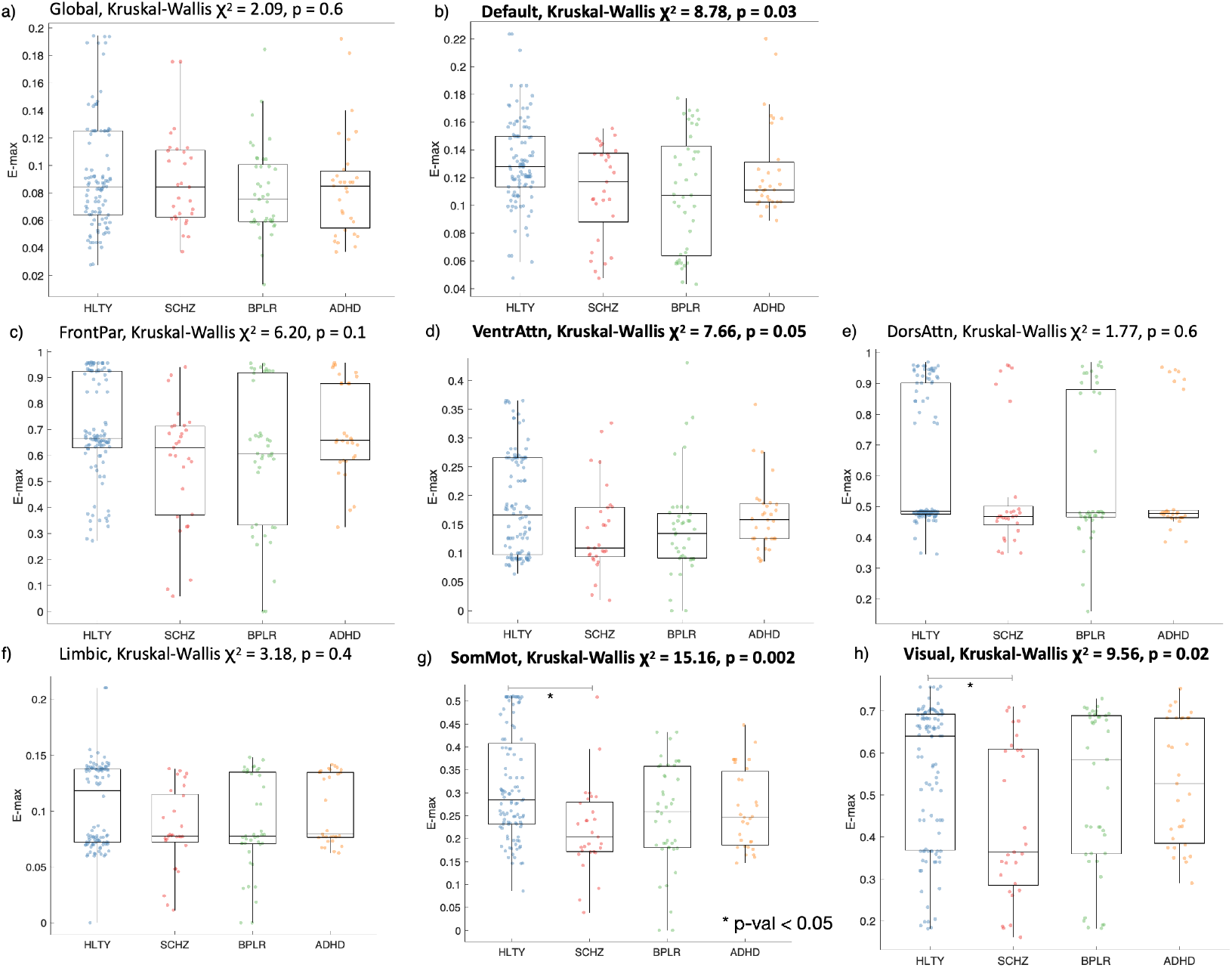
Energy gap metrics: Boxplots showing group E-max distributions when values are a) global, or restricted to each of the seven resting state networks, which are the b) default mode, c) frontal parietal, d) ventral attention, e) dorsal attention, f) limbic, g) somatomotor, and h) visual. The subpanel titles show the Kruskal-Wallis test results for group comparisons after controlling for age, sex, site, and average FD (Bolded ones have a significant overall group effect, p < 0.05). For comparisons with a significant overall group effect, post hoc ranksum tests were done between each group pair. Significant paired differences after Tukey correction (* p-val < 0.05) are marked by a line and star. There are main group effects for the default mode, ventral attention, somatomotor, and visual networks. Post hoc comparisons show larger E-max in the HLTY group than SCHZ group for the somatomotor and visual networks.

### Network specific energy gap differs across groups

As an exploratory analysis, the DK atlas parcels were assigned into the 7 canonical networks (Supplemental Fig. 10a). Significant group differences in terms of E-max and E-mean were present for certain networks, with changes not necessarily in the same direction. E-max comparison results for each network are shown in Figure 4, while those for E-mean are shown in Supplemental Fig. 11.

Significant overall group differences in terms of E-max were found for: 1) the default mode network (𝒳^2^ = 8.78, p = 0.03), which did not show any significant paired differences after correction; 2) the ventral attention network, which did not show any significant paired differences after correction; 3) the somatomotor network (𝒳^2^ = 15.16, p = 0.002), which had a significantly higher value for the HLTY group than the SCHZ group, and 4) the visual network (𝒳^2^ = 9.56, p = 0.02), which also had a significantly higher value for the HLTY group than the SCHZ group.

No significant group differences in terms of E-mean were found globally or for any resting state networks.

### Effect of group specific SC on fitting results

To examine how our study findings are affected by the choice of group specific SC, we repeated our analyses by fitting cross-attractor coordination matrices generated with the healthy group SC for all the groups. Results showed similar trends in the group comparisons but with less significant findings. For example, while global coupling (i.e., *G*) still had a significant group effect, there was no longer a significant paired difference between HLTY and SCHZ (Supplemental Fig. 12b). The preservation of some effects, but not others support the need for group specific SC, which has a nonlinear effect on the model fitting. The results with HLTY SC are shown in Supplemental Fig. 12, 13, and 14, which corresponds to Figure 3, Figure 4, and Supplemental Fig. 11 respectively.

### Examining correlation with clinical symptoms

Given the significant group differences in global coupling and w_EE_/w_EI_ ratio, we sought to determine if they were linked with specific symptoms while controlling for age, sex, site (scanner), and average FD. For ADHD participants, we correlated with their hyperactivity and attention total scores on the Adult ADHD Clinical Diagnostic Scale (ACDS). For BPLR participants, we correlated with their total score on the Young’s Mania Rating Scale (YMRS) and their total score on the 17-items Hamilton Depression Rating Scale (HAMD-17). For SCHZ participants, we correlated the parameters with their total scores on the Scale for the Assessment of Positive Symptoms (SAPS) and the Scale for the Assessment of Negative Symptoms (SANS). Through our exploratory analysis, we found that the parameter *G* was inversely correlated with the total score for SAPS (Rho = -0.51, p = 0.009). No other correlations were significant.

## Discussion

Our results demonstrated that psychiatric disorders might be characterized by disturbances in the brain’s attractor landscape described by our BNM model. Specifically, significant group effects were found for the global coupling parameter and the E/I ratio between local excitation and inhibition parameters. All patient groups had an average global coupling value lower than that of healthy controls, with the schizophrenia group being significantly different. The E/I ratio was also different across the groups, with the schizophrenia group having a significantly larger value than the ADHD group. Further insight was revealed by comparing measures of energy gap associated with the optimal individual parameters. Specifically, there were group effects for the maximum energy gap constrained to the default mode, ventral attention, somatomotor, and visual networks.

### Model fitting performance across psychiatric populations

The level of fit between the cross-attractor coordination matrix and the real RSFC was similar across the groups. The level of correlation is comparable to reported values from previous BNM studies (Gustavo Deco et al., 2013; Iravani et al., 2021; Park et al., 2021; Wang et al., 2019), suggesting that the model fitting procedure works well for patient populations.

### Global coupling differences across psychiatric populations

There was an overall group effect for the global coupling *G*, with the healthy group having the highest value (*G* = 3.0), followed by the ADHD (*G* = 2.9) and bipolar (*G* = 2.6) groups, and with the schizophrenia group having the lowest (*G* = 2.4). Paired comparisons show that *G* was significantly less for schizophrenia patients than healthy controls. The finding suggests that psychopathology may be partly due to changes in the global coupling modulating connections between all regions. A recent study argued that global coupling could be considered a factor for operationalizing the balance between local and global influences on a brain region, such that a decrease in global coupling could result in less (or more) global (or local) influence (Klein et al., 2021). Here we found that a decrease in global coupling for schizophrenia patients was correlated with more positive symptoms. This is consistent with the disconnection hypothesis (Friston et al., 2016) for schizophrenia, which attributes symptoms to a disruption of normal large-scale brain network dynamics.

### Differences in excitation-inhibition ratio across populations

Our results show that the E/I ratio for healthy individuals as captured by average *w*_*EE*_/*w*_*EI*_ ratio was 1.02 (0.72). Our previous work had shown that a model with a fixed *w*_*EE*_/*w*_*EI*_ ratio of 2 was still able to significantly predict the real RSFC (Zhang et al., 2022), which suggests that parts of the RSFC variance may be insensitive to changes in the local parameters. Nevertheless, the improvement in prediction based on our individually fitted parameters points to the importance of capturing individual variability.

The attractor landscape for the optimal individual configurations was dominated by stable spirals or damped oscillations (see Supplemental Fig. 15 to 22). While the reasoning for this observation may require further theoretical exploration, one possibility is that the ongoing activity of the resting state brain is a summation of damped oscillatory processes (Evertz et al., 2022), which can arise from spontaneous transitions between the available stable spiral attractors.

Our results show that schizophrenia patients have the highest E/I ratio as captured by an average *w*_*EE*_/*w*_*EI*_ ratio of 1.30 (0.78), while ADHD patients have the lowest E/I ratio, with an average *w*_*EE*_/*w*_*EI*_ ratio of 0.89 (0.77). Abnormal E/I balance has been consistently linked to psychiatric disorders (Sohal and Rubenstein, 2019). In the case of schizophrenia patients, an abnormally high E/I ratio can be attributed to deficits in GABA mediated inhibitory neurotransmission (Benes and Berretta, 2001; Marín, 2012). Likewise, abnormalities in E/I balance, both during development and persisting into adulthood, have been attributed to ADHD patients as a neurobiological cause (Mamiya et al., 2021). Indeed, an abnormally low E/I ratio may be a result of deficits in glutamate based excitatory neurotransmission (Cheng et al., 2017; Dramsdahl et al., 2011).

### Implications of differences in energy gap metrics

In terms of maximum energy gaps, there was an overall group effect for the default mode network. Relative to the healthy controls, all patient groups had lower maximum energy gaps, with the schizophrenia group having the largest abnormality. This finding is supported by studies showing default mode network abnormalities in schizophrenia patients (Bluhm et al., 2007; Ongür et al., 2010; Whitfield-Gabrieli and Ford, 2012). Since maximum energy gaps are essentially gaps between potential sub-repertoires of attractors, lower values may result in excessive transitions within a specific network (Al Zoubi et al., 2019; da Cruz et al., 2020).

E-max was also significantly lower for schizophrenia subjects relative to healthy controls in the somatomotor and visual networks. Similar to the case of the default mode network, a lower E-max for the somatomotor and visual networks may cause abnormal sensory activation (Li et al., 2017). Findings of abnormalities in the somatomotor network agrees with one recent study focusing on the RSFC of the same dataset, which showed a link between psychopathology across the disorders and functional connectivity of the somatomotor network (Kebets et al., 2019). Likewise, another study examining task-based FC of the same dataset showed group differences in regions part of the frontal parietal and somatomotor networks (Barron et al., 2021).

### Limitations and future directions

While the results of this study showed the promise of our modeling approach for understanding psychiatric disorders, there are several limitations. First, the sample size is small, especially for the patient groups. Therefore, replication of the results in a larger and independent sample is necessary. Nevertheless, given that the purpose of this study was to demonstrate the utility of the cross-attractor modeling approach in patient populations, the immediate goal was achieved. Second, the quality of the DWI and rsfMRI scans was poor, especially relative to the quality of HCP scans, which is likely why the model fitness values for the groups were slightly lower than reported for the sample of HCP subjects in our previous study (Zhang et al., 2022). While our model was able to draw helpful conclusions from the clinical grade scans, refinement of the model will benefit from having higher quality scans for patient populations. Indeed, although using a group-specific SC instead of one for all groups allowed for preserving group variations in SC in the subsequent analysis, individual SC might better account for individual differences in the structural connectome and should be used if possible. Third, while the parcellation we used was a good compromise between computational run time and biological realism, higher resolved parcellations will be explored in our future studies. Moreover, we plan to examine dynamic FC in addition to regular FC for model fitting and will evaluate how the fitted model parameters differ across groups. Dynamic FC has been shown to capture faster changing temporal dynamics (Allen et al., 2014; Cabral et al., 2017; Zhang et al., 2023). Therefore, examining changes in model parameters fitted to dynamic FC will likely reveal more insights into the pathophysiology of the patient groups. Likewise, the resolution of our parameter search space may limit the accuracy of the individual fitting results. Still, it was selected with considerations for computation time in this proof-of-concept study. Finally, the model did not allow the local parameters to vary between regions. The assumption of uniform configurations across the brain had been made in most BNM studies due to computational limitations and the risk of overfitting (Deco et al., 2013; Schirner et al., 2018). Future studies with optimization approaches built for high dimensional fitting problems, such as the evolutionary optimization (Maile et al., 2019; Miikkulainen, 2021), may help to overcome this challenge.

## Data and code availability statement

The LA5c dataset was available through the OpenNeuro website, with further details presented elsewhere (Poldrack et al., 2016). The code for the w3c model and for computing the cross-attractor coordination is available online. Further details of the analysis pertaining to the LA5c dataset is available from the authors upon reasonable request.

## Ethics statement

All participants gave informed consent and were either healthy (HLTY) or had a clinical diagnosis of schizophrenia (SCHZ), bipolar disorder (BPLR), or attention deficit hyperactivity disorder (ADHD).

