## Supplementary Information for "Cross-attractor modeling of resting-state functional connectivity in psychiatric disorders"

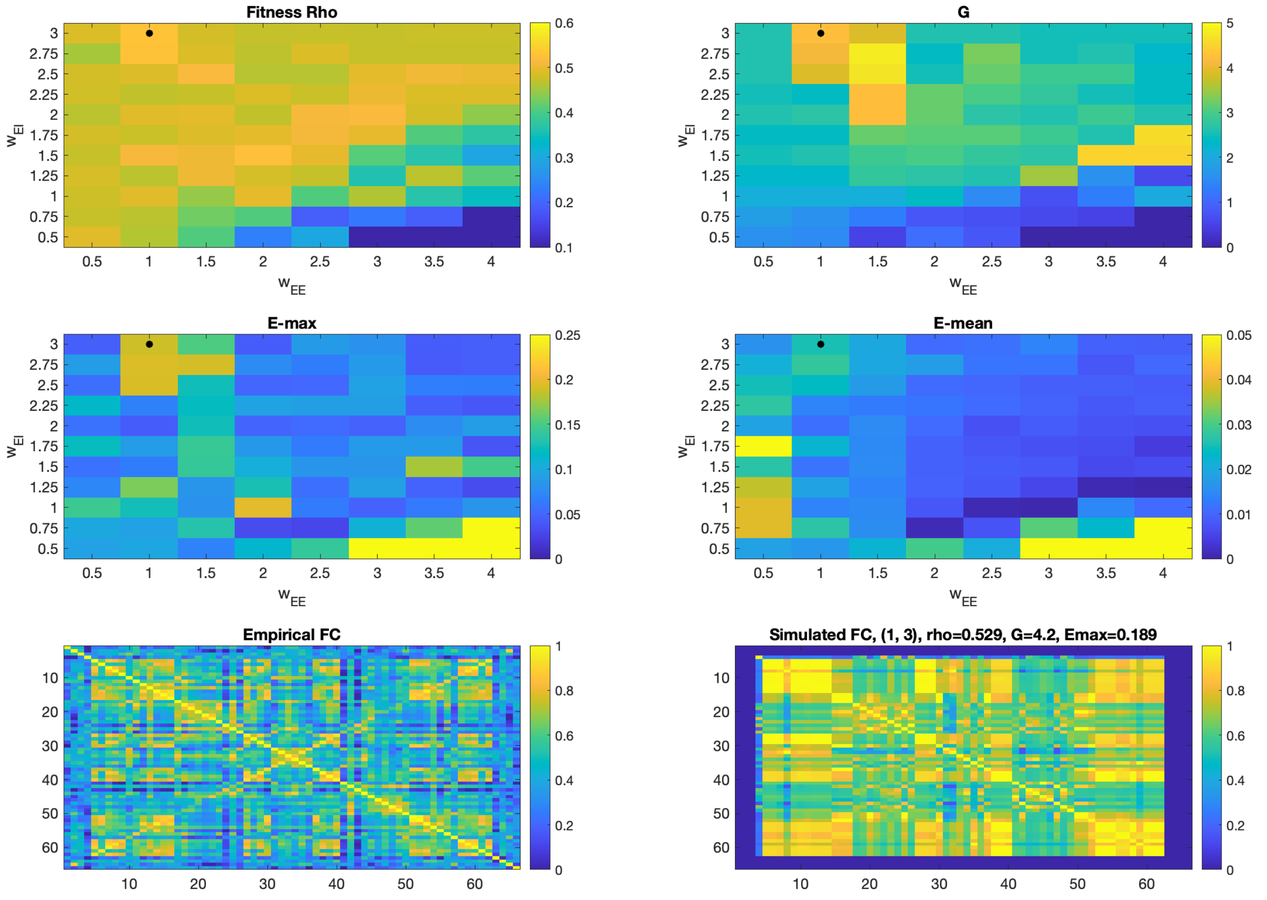


**Supplemental Fig. 1:** Example individual fitting result for a HLTY participant (sub-10517)

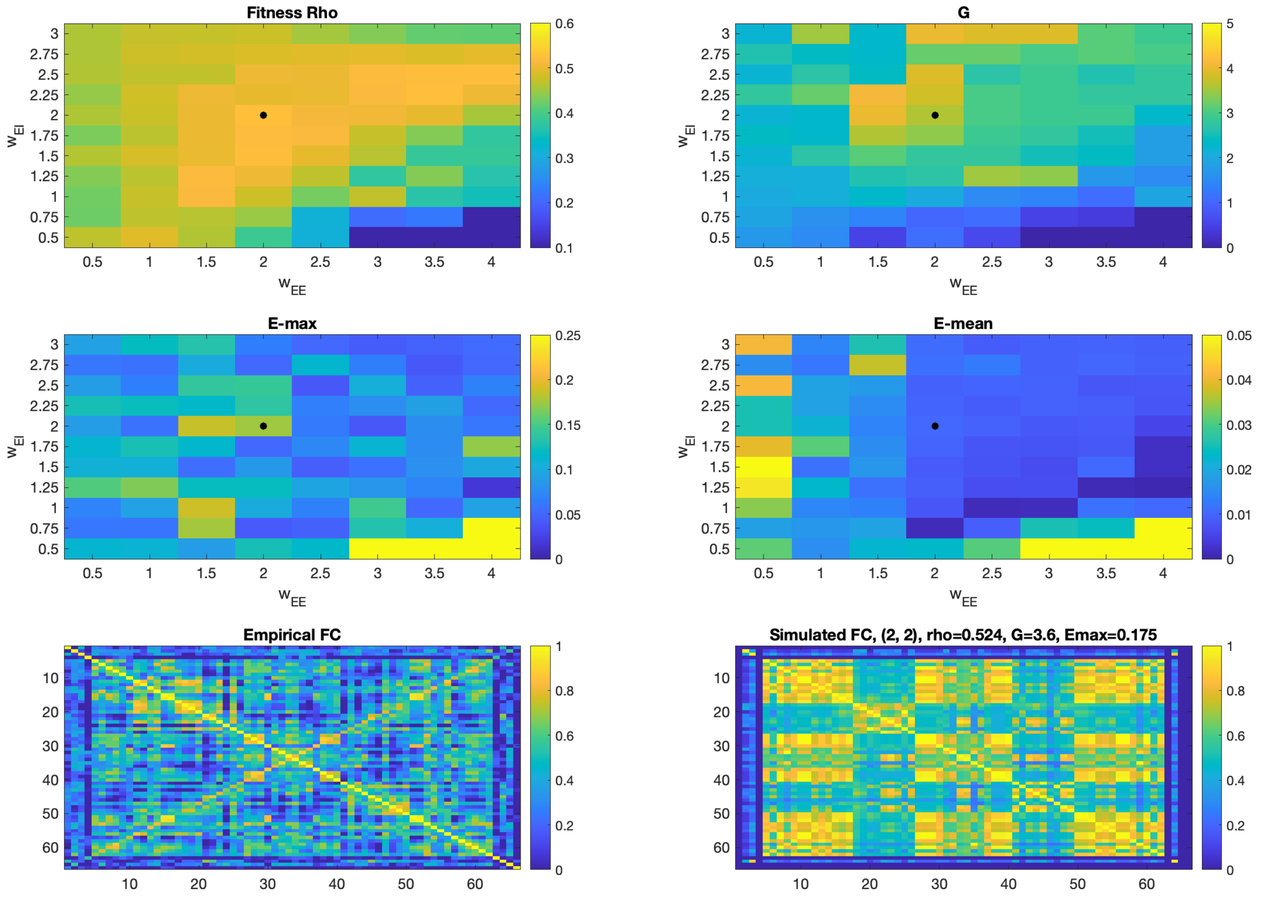


**Supplemental Fig. 2:** Example individual fitting result for a SCHZ participant (sub-50055)

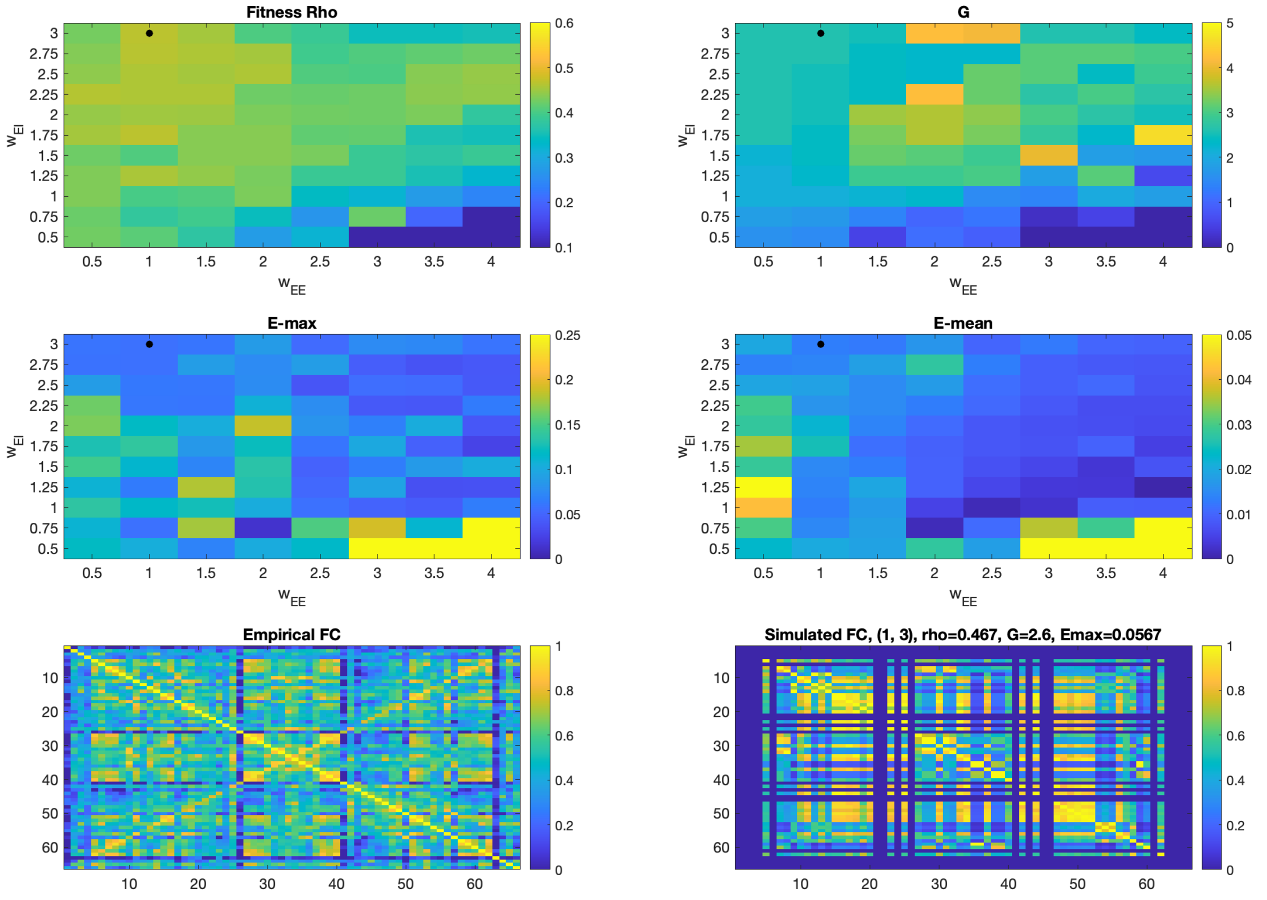


**Supplemental Fig. 3:** Example individual fitting result for a BPLR participant (sub-60079)

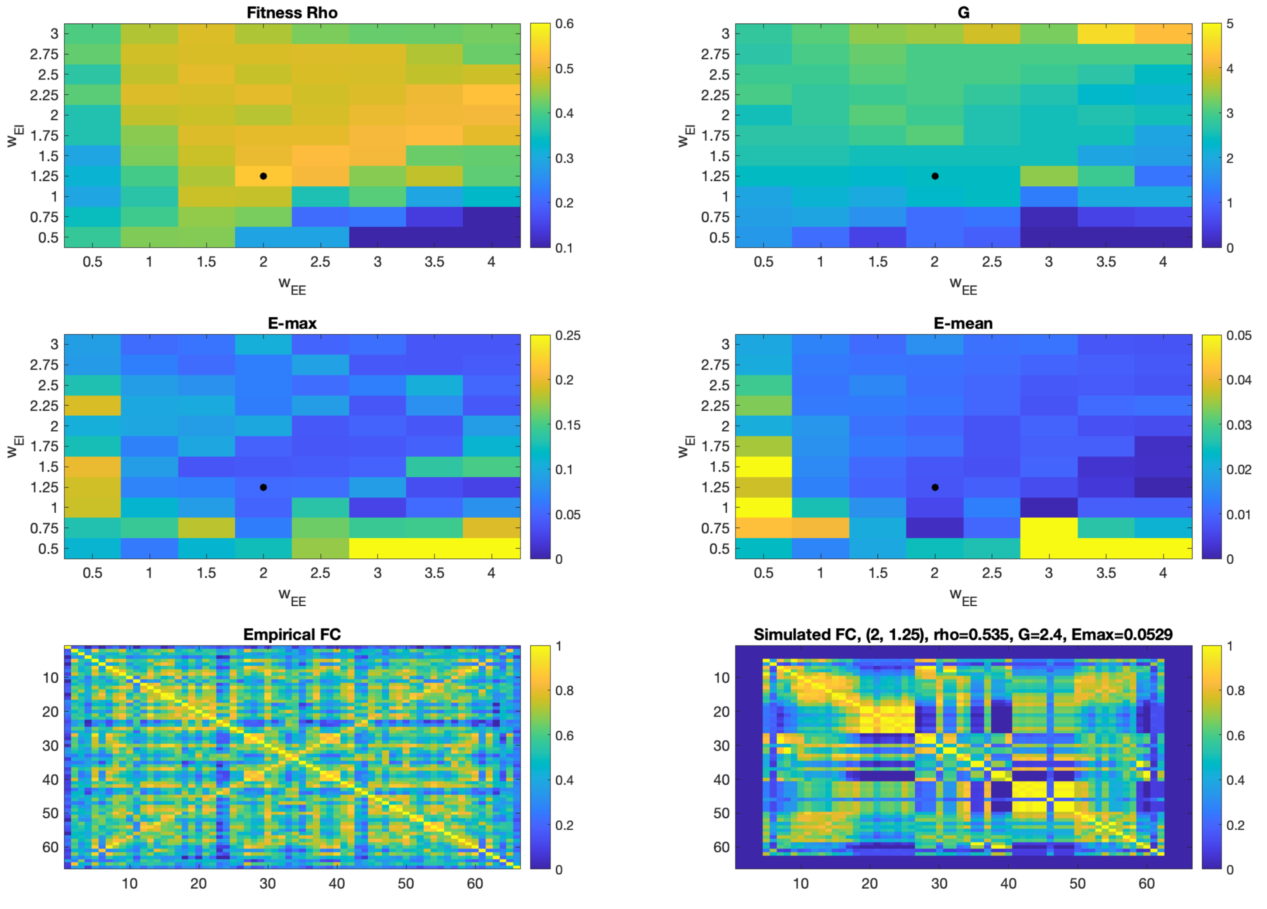


**Supplemental Fig. 4:** Example individual fitting result for an ADHD participant (sub-70051)

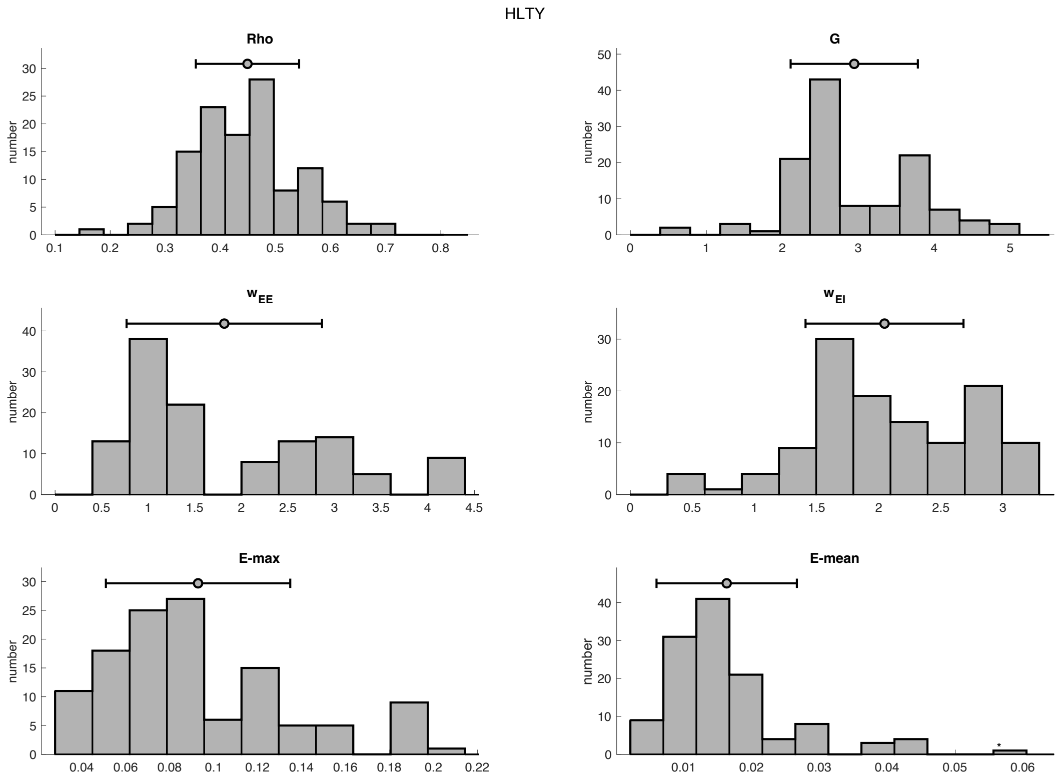


**Supplemental Fig. 5:** Distribution of model parameters and metrics for healthy (HLTY) participants

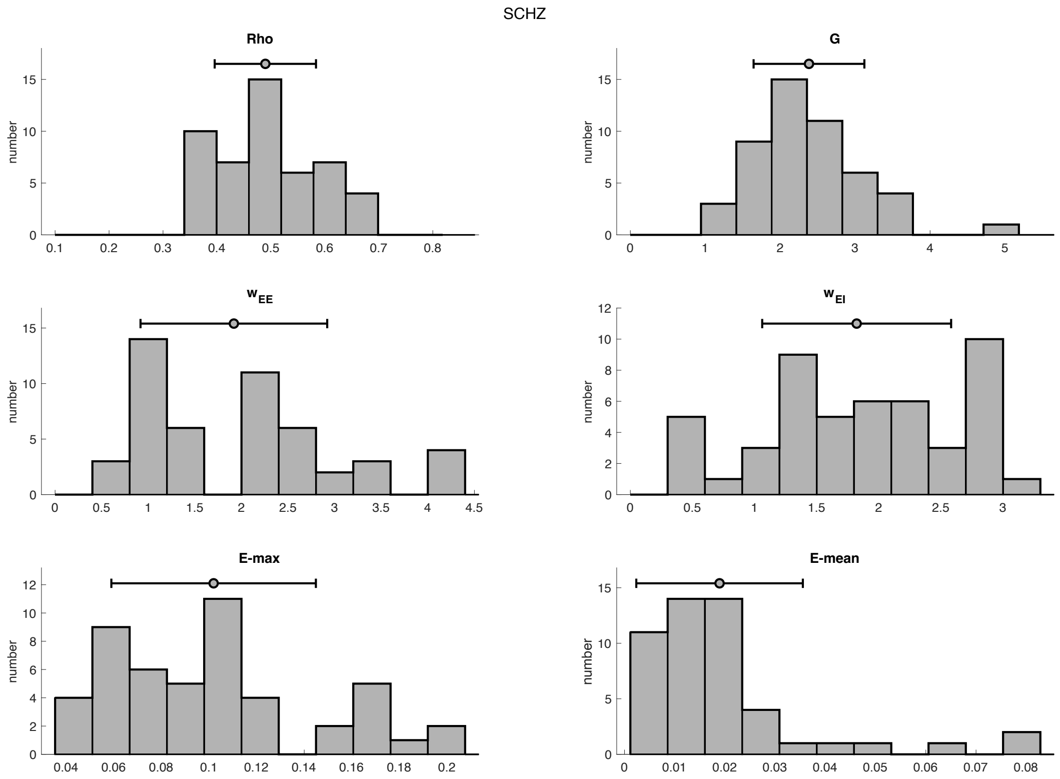


**Supplemental Fig. 6:** Distribution of model parameters and metrics for schizophrenia (SCHZ) participants

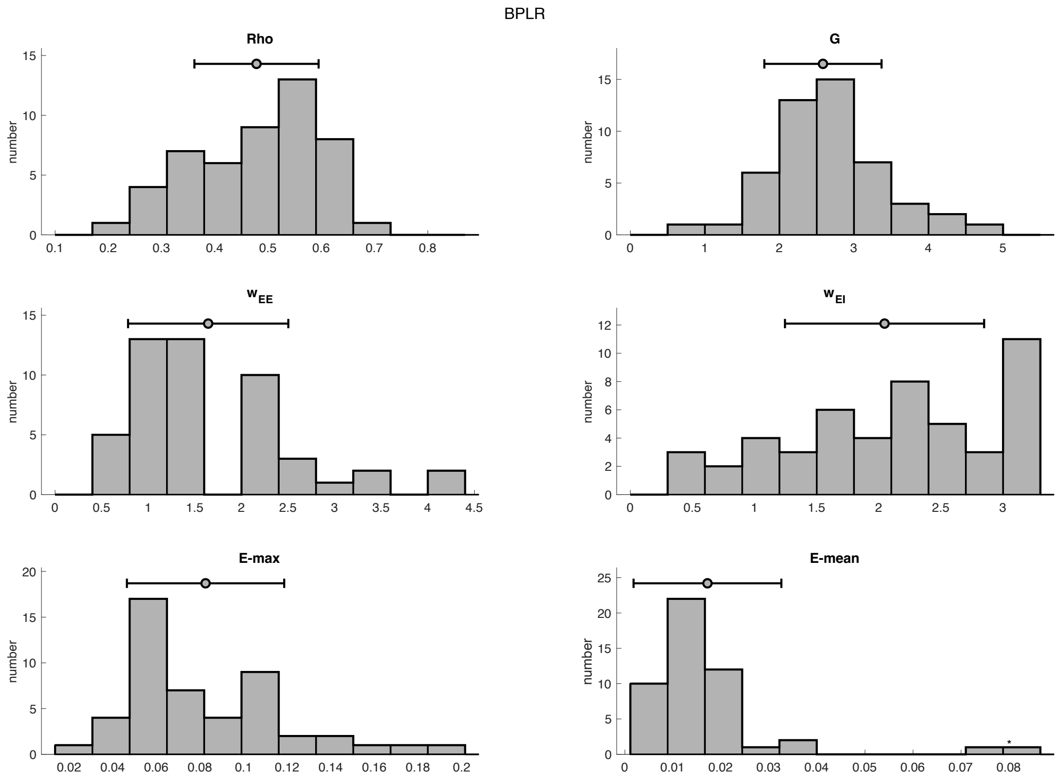


**Supplemental Fig. 7:** Distribution of model parameters and metrics for bipolar (BPLR) participants

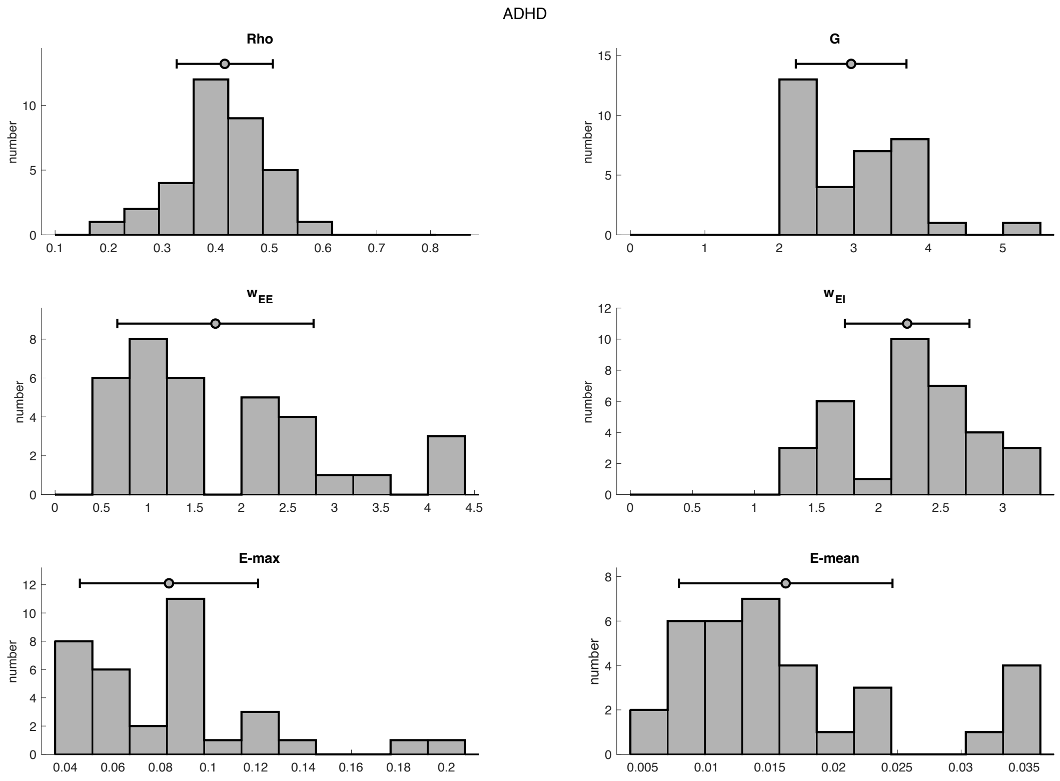


**Supplemental Fig. 8:** Distribution of model parameters and metrics for ADHD participants

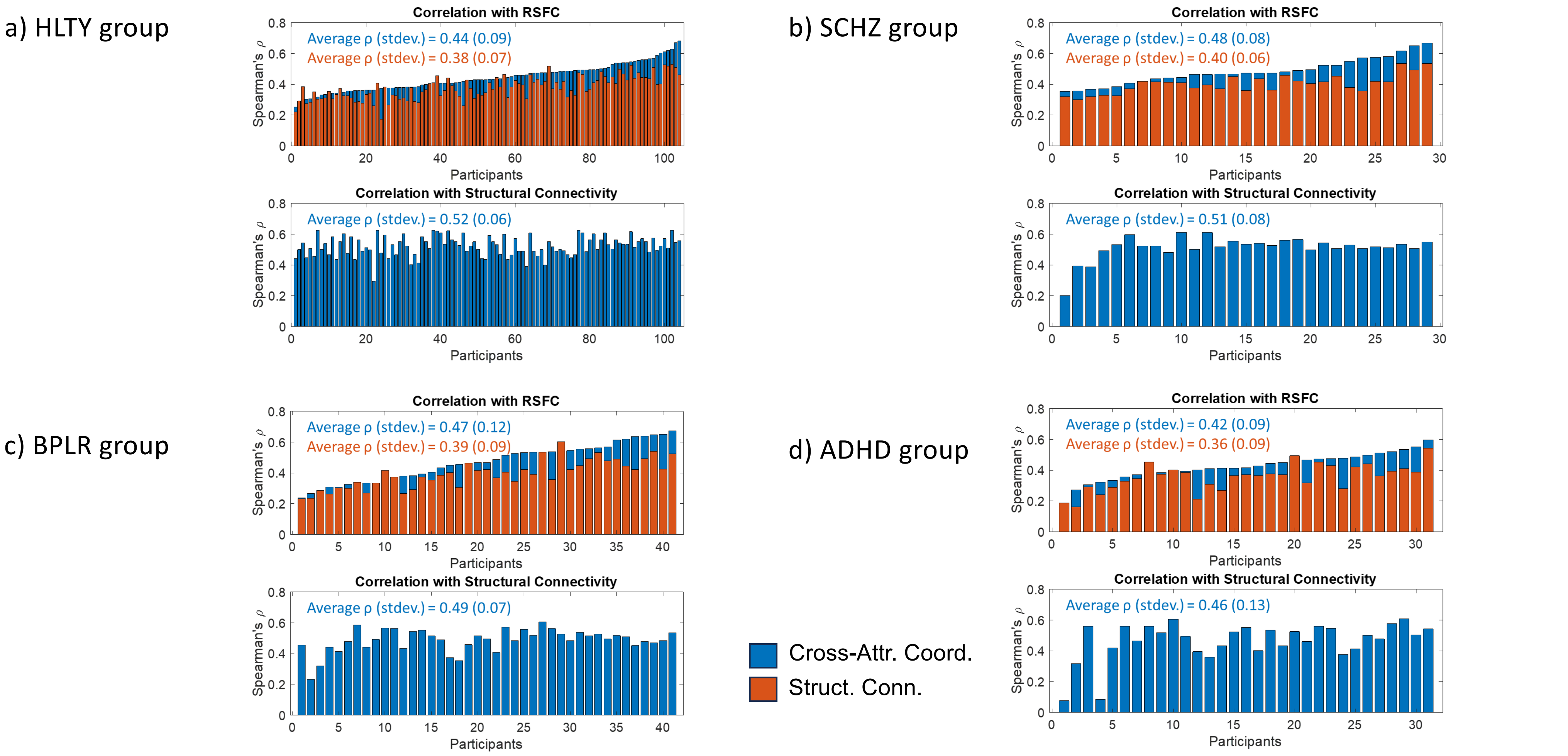


**Supplemental Fig. 9:** Paired correlations between individual RSFC, cross-attractor coordination optimally fitted to the individual RSFC, and group SC for a) HLTY, b) SCHZ, c) BPLR, and d) ADHD participants. The participant values are sorted in ascending order for the correlation between cross-attractor coordination and individual RSFC.

a)

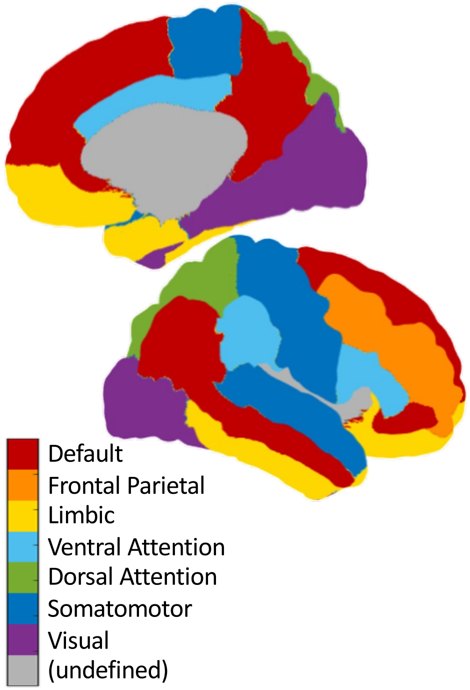


b)

| | Right Hemisphere | | Left Hemisphere | |
|---|---|---|---|---|
| **Parcel Name** | **Network** | **Index** | **Network** | **Index** |
| entorhinal | Limbic | 1 | Limbic | 66 |
| parahippocampal | Visual | 2 | Default | 65 |
| temporalpole | Limbic | 3 | Limbic | 64 |
| frontalpole | Limbic | 4 | Limbic | 63 |
| fusiform | Visual | 5 | Visual | 62 |
| transversetemporal | SomMot | 6 | SomMot | 61 |
| lateraloccipital | Visual | 7 | Visual | 60 |
| superiorparietal | DorsAttn | 8 | DorsAttn | 59 |
| inferiortemporal | Limbic | 9 | Limbic | 58 |
| inferiorparietal | Default | 10 | Default | 57 |
| supramarginal | VentAttn | 11 | VentAttn | 56 |
| bankssts | SomMot | 12 | Default | 55 |
| middletemporal | Default | 13 | Default | 54 |
| superiortemporal | SomMot | 14 | SomMot | 53 |
| postcentral | SomMot | 15 | SomMot | 52 |
| precentral | SomMot | 16 | SomMot | 51 |
| caudalmiddlefrontal | FrontPar | 17 | Default | 50 |
| parsopercularis | VentAttn | 18 | VentAttn | 49 |
| parstriangularis | VentAttn | 19 | Default | 48 |
| rostralmiddlefrontal | FrontPar | 20 | FrontPar | 47 |
| parsorbitalis | Default | 21 | Default | 46 |
| lateralorbitofrontal | Limbic | 22 | Limbic | 45 |
| caudalanteriorcingulate | VentAttn | 23 | VentAttn | 44 |
| rostralanteriorcingulate | Default | 24 | Default | 43 |
| superiorfrontal | Default | 25 | Default | 42 |
| medialorbitofrontal | Limbic | 26 | Limbic | 41 |
| lingual | Visual | 27 | Visual | 40 |
| pericalcarine | Visual | 28 | Visual | 39 |
| cuneus | Visual | 29 | Visual | 38 |
| paracentral | SomMot | 30 | SomMot | 37 |
| isthmuscingulate | Default | 31 | Default | 36 |
| precuneus | Default | 32 | Default | 35 |
| posteriorcingulate | VentAttn | 33 | VentAttn | 34 |

**Supplemental Fig. 10:** For comparing E-max and E-mean, parcels are assigned into 7 canonical networks as shown in a), which correspond to table in b)

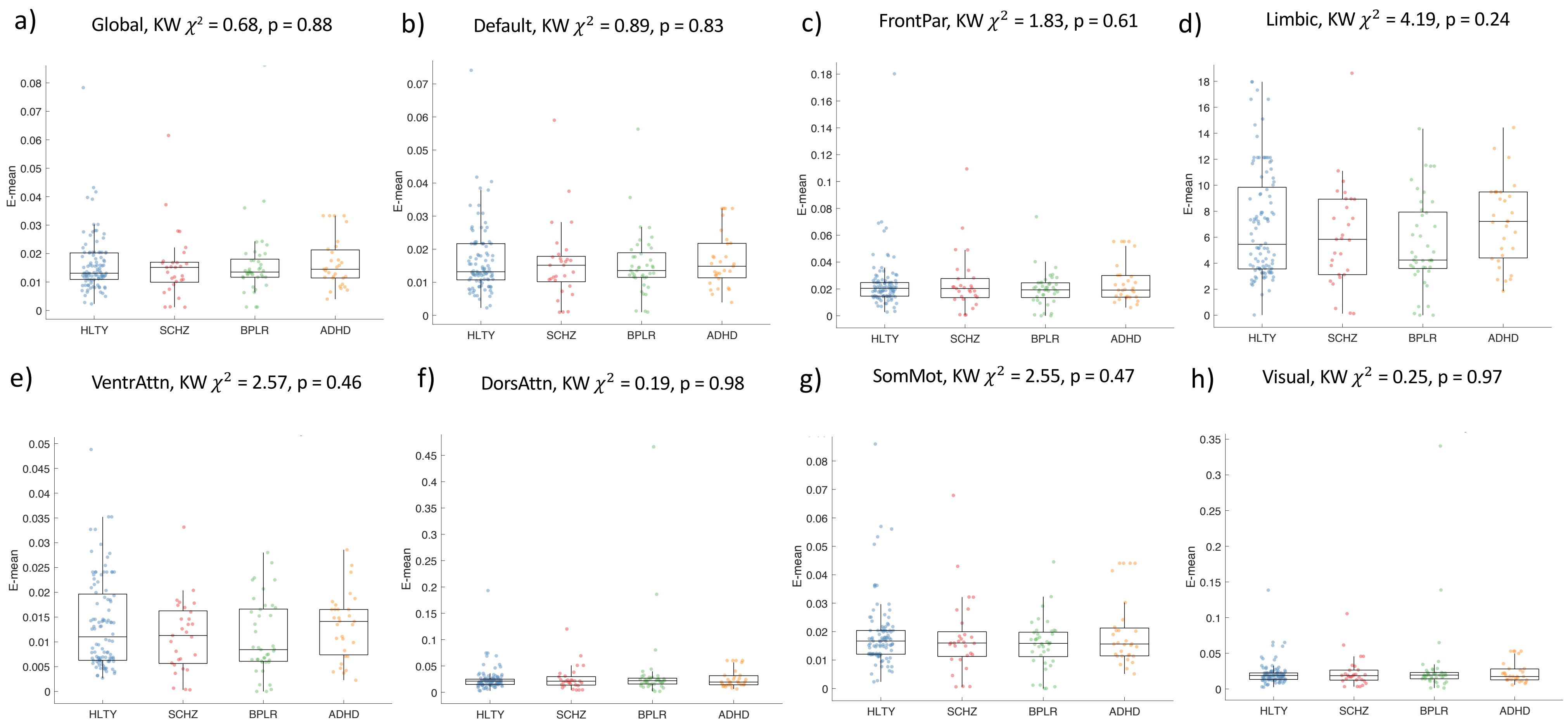


**Supplemental Fig. 11:** For comparing E-mean, parcels are assigned into 7 canonical networks as shown in Supplemental Fig. 9 a). Boxplots show the group distributions of the a) global E-mean, while b) to h) shows the group distributions of E-mean for each network. The subtitle shows the Kruskal-Wallis (KW) test results for group comparisons after controlling for age, sex, site, and average FD.

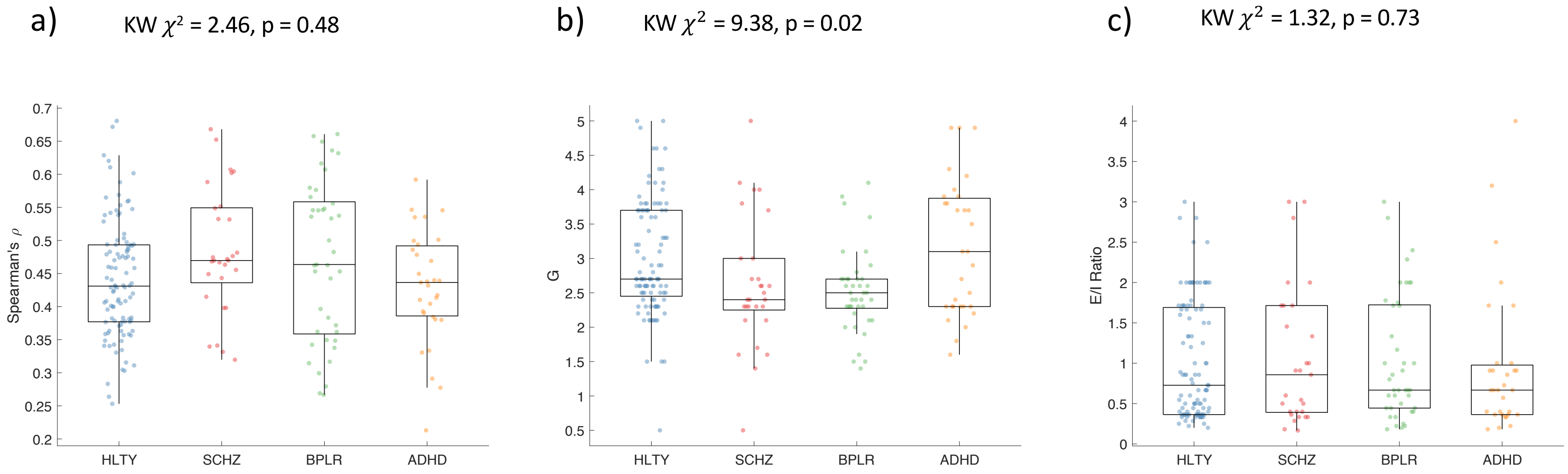


**Supplemental Fig. 12:** Repeat of Figure 2 with HLTY simulation results fitted for all groups. Model parameter fitting: Boxplots showing the distribution across group participants for a) model fitness (Spearman's ρ), b) global coupling (G) value, and c) excitation to inhibition ratio ($w_{EE}/w_{EI}$). The Kruskal-Wallis (KW) test results are shown for group comparisons after controlling for age, sex, site, and average FD.

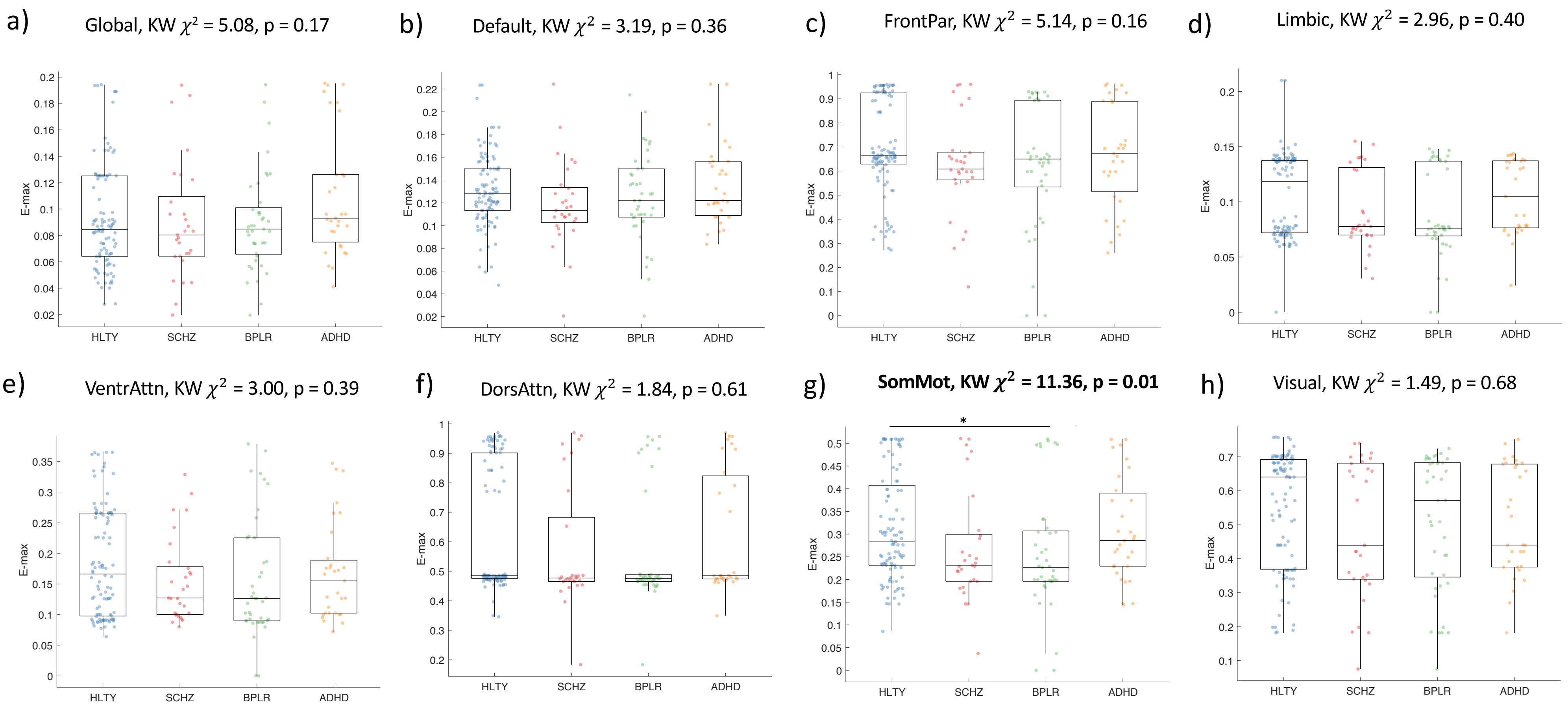


**Supplemental Fig. 13:** Repeat of Figure 3 with HLTY simulation results fitted for all groups. For comparing E-max, parcels are assigned into 7 canonical networks as shown in Supplemental Fig. 9 a). Boxplots show the group distributions of the a) global E-mean, while b) to h) shows the group distributions of E-max for each network. The subtitle shows the Kruskal-Wallis (KW) test results for group comparisons after controlling for age, sex, site, and average FD (Bolded ones have a significant overall group effect, $p < 0.05$) .

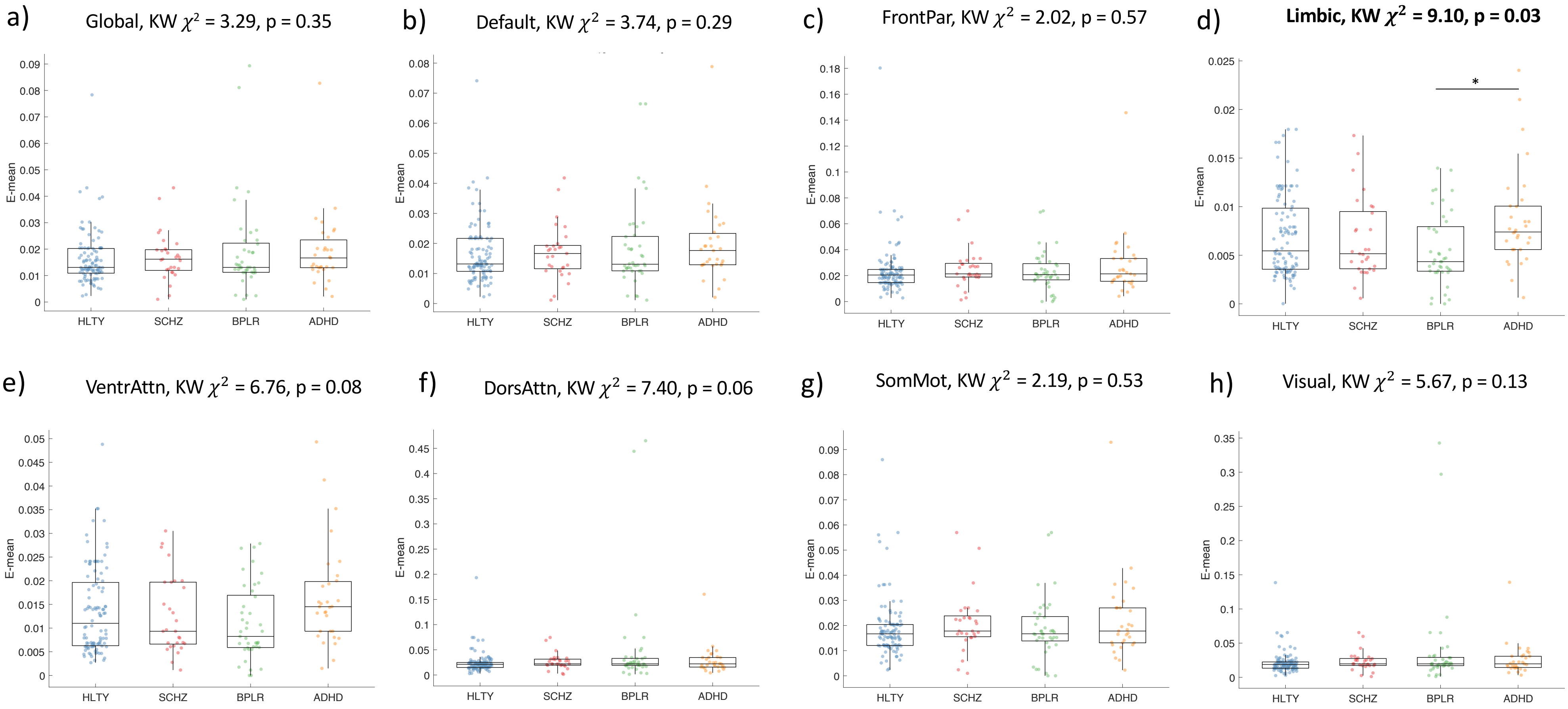


**Supplemental Fig. 14:** Repeat of Supplemental Figure 10 with HLTY simulation results fitted for all groups. For comparing E-mean, parcels are assigned into 7 canonical networks as shown in Supplemental Fig. 9 a). Boxplots show the group distributions of the a) global E-mean, while b) to h) shows the group distributions of E-mean. The subtitle shows the Kruskal-Wallis (KW) test results for group comparisons after controlling for age, sex, site, and average FD (Bolded ones have a significant overall group effect, $p < 0.05$) .

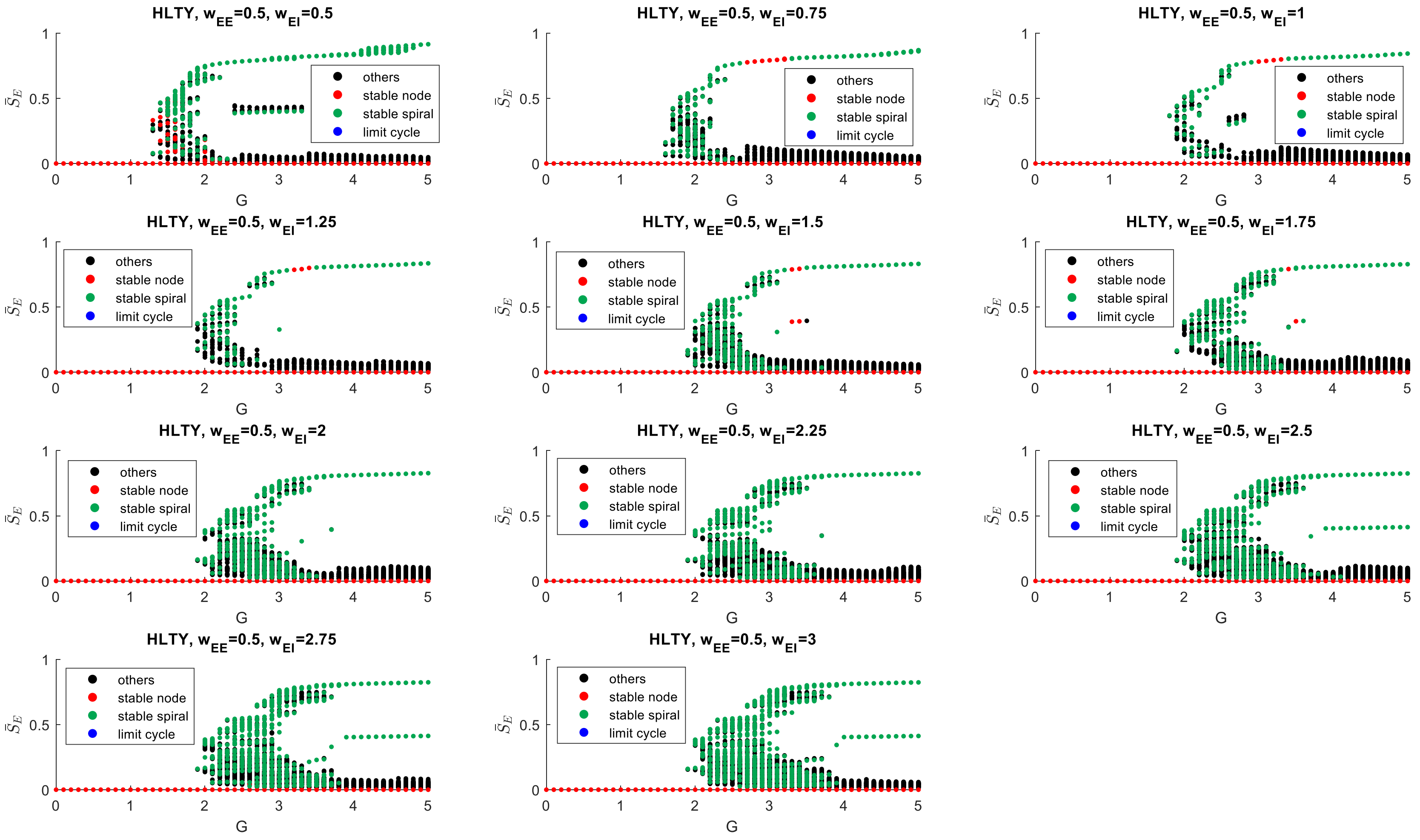


**Supplemental Fig. 15:** Population-level bifurcation plots for healthy controls ($W_{EE}$ = 0.5)

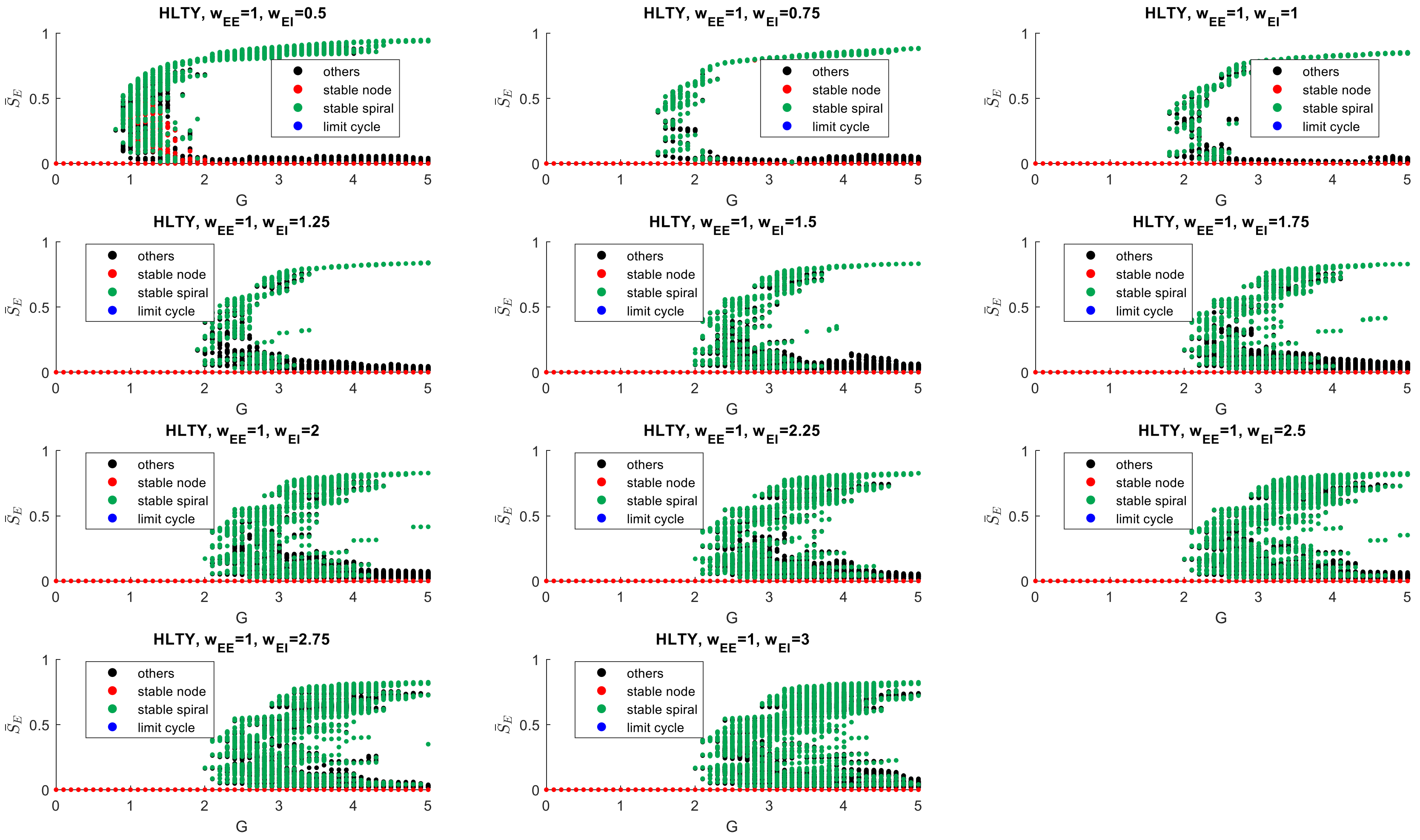


**Supplemental Fig. 16:** Population-level bifurcation plots for healthy controls ($W_{EE}$ = 1.0)

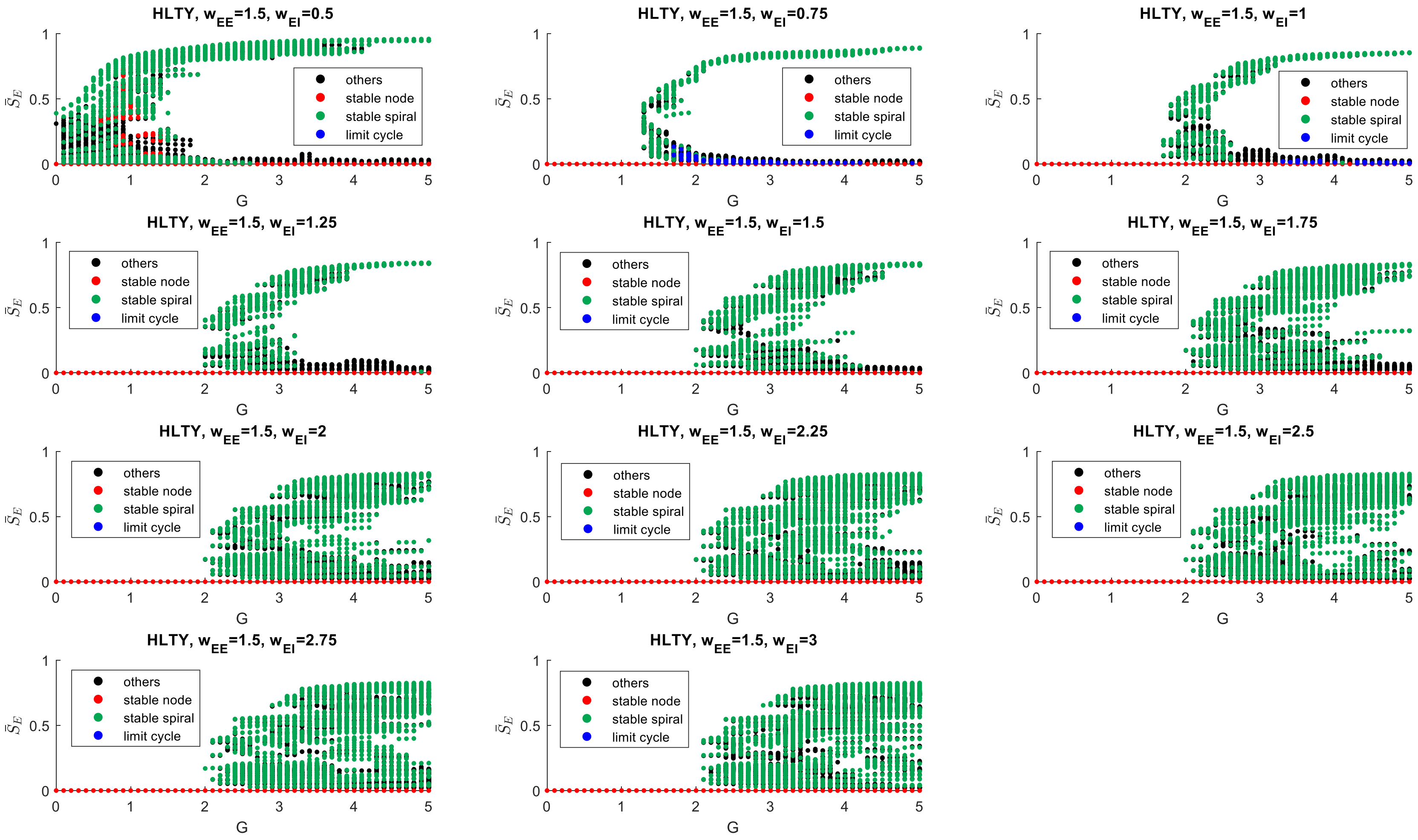


**Supplemental Fig. 17:** Population-level bifurcation plots for healthy controls ($W_{EE}$ = 1.5)

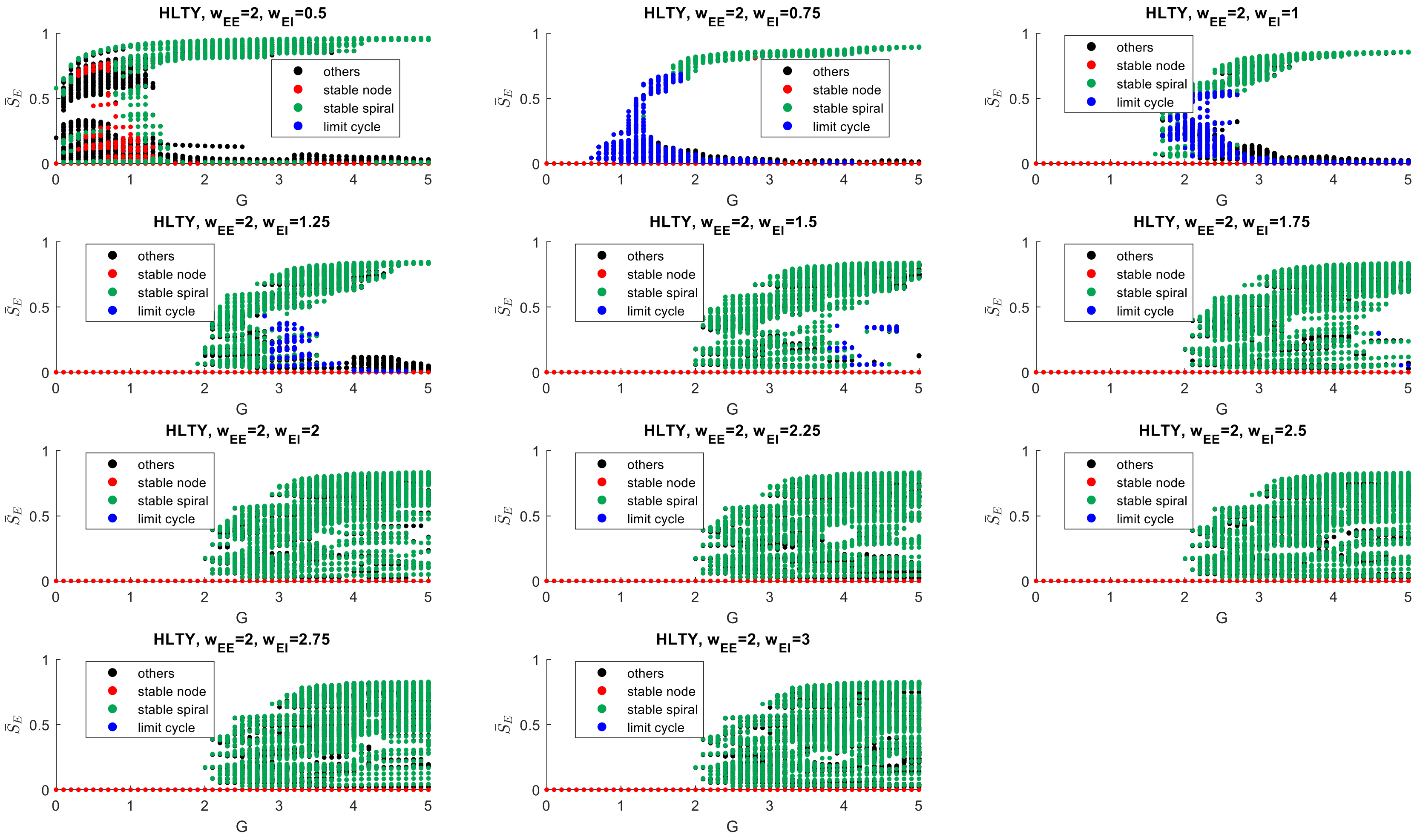


**Supplemental Fig. 18:** Population-level bifurcation plots for healthy controls ($W_{EE}$ = 2.0)

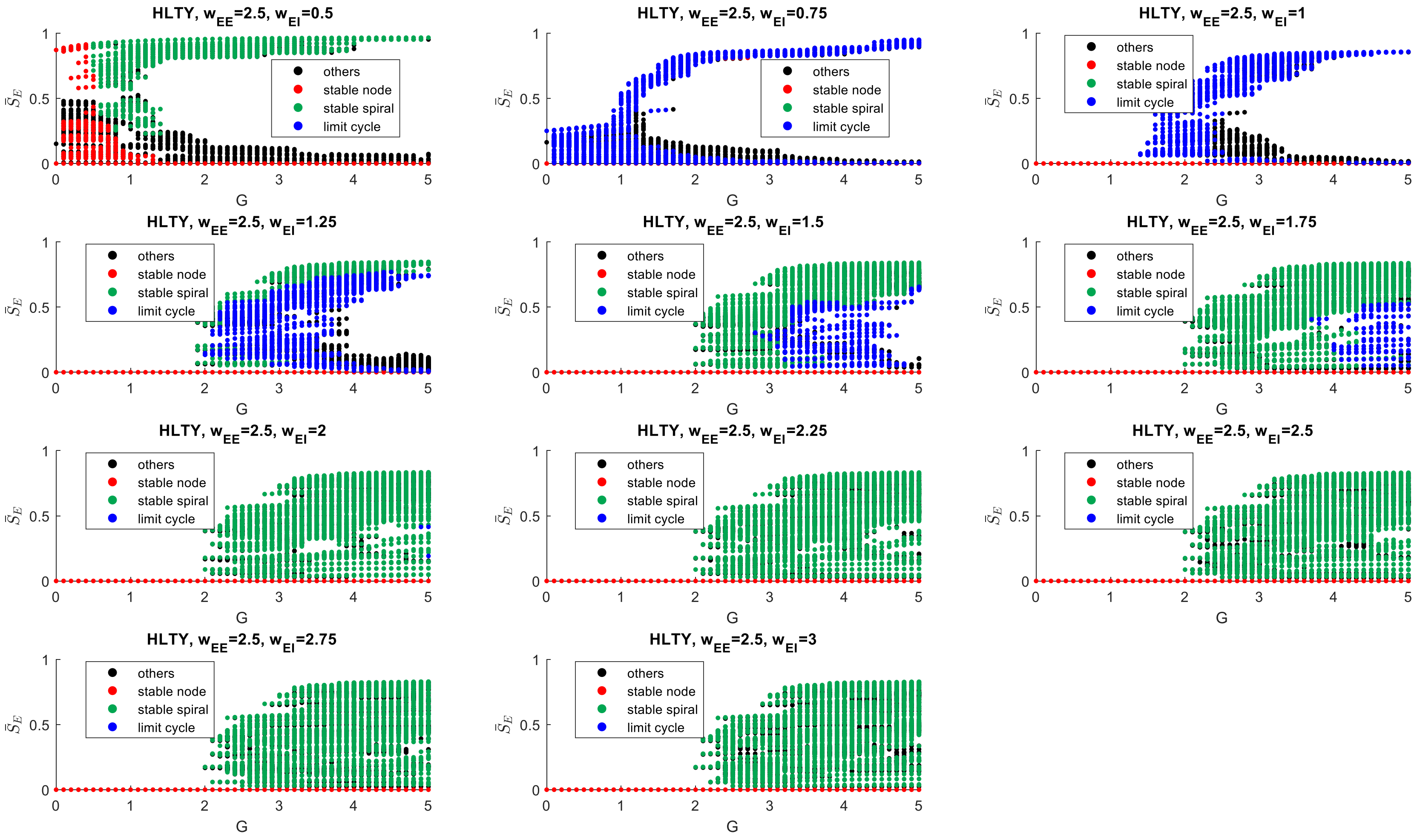


**Supplemental Fig. 19:** Population-level bifurcation plots for healthy controls ($W_{EE}$ = 2.5)

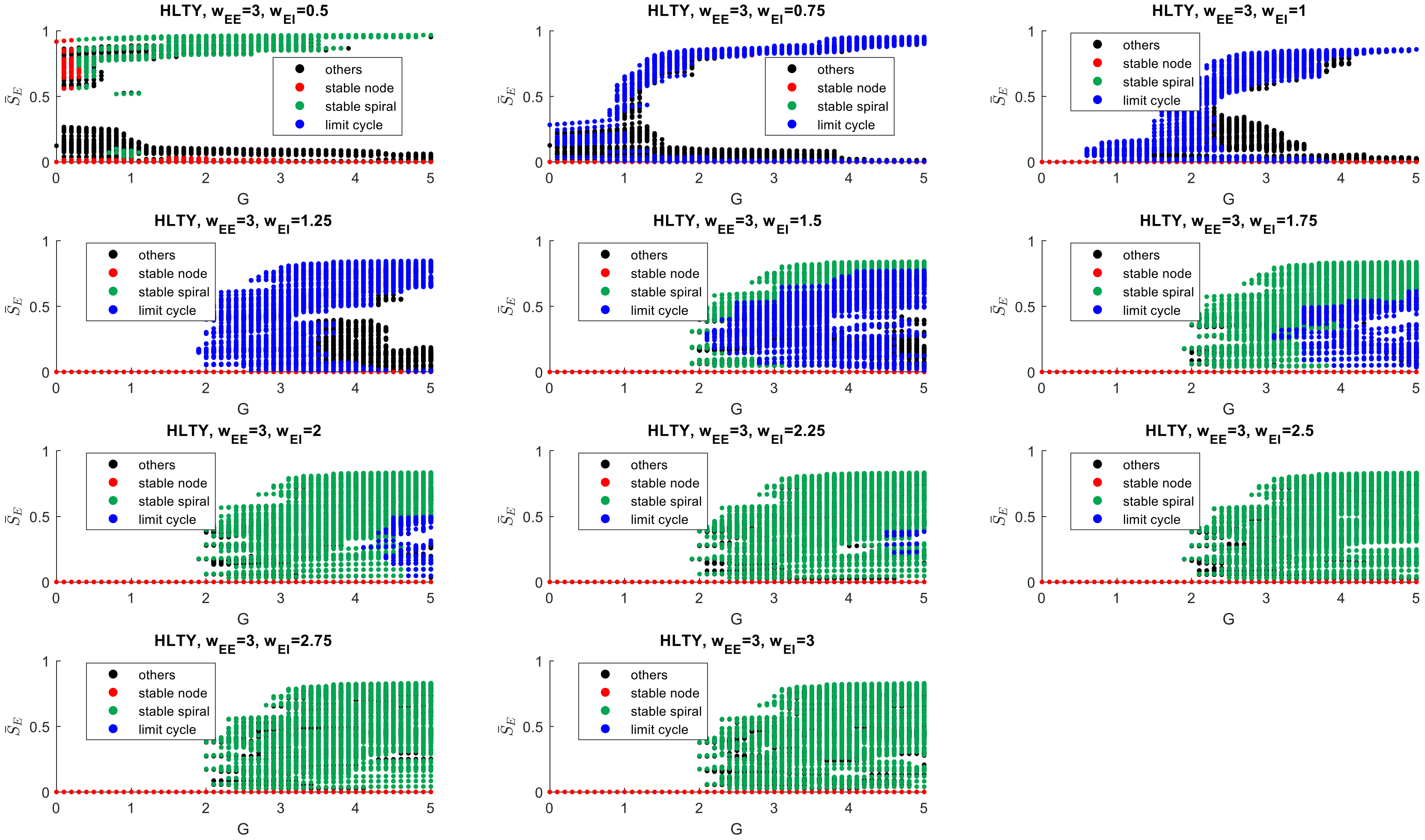


**Supplemental Fig. 20:** Population-level bifurcation plots for healthy controls ($W_{EE}$ = 3.0)

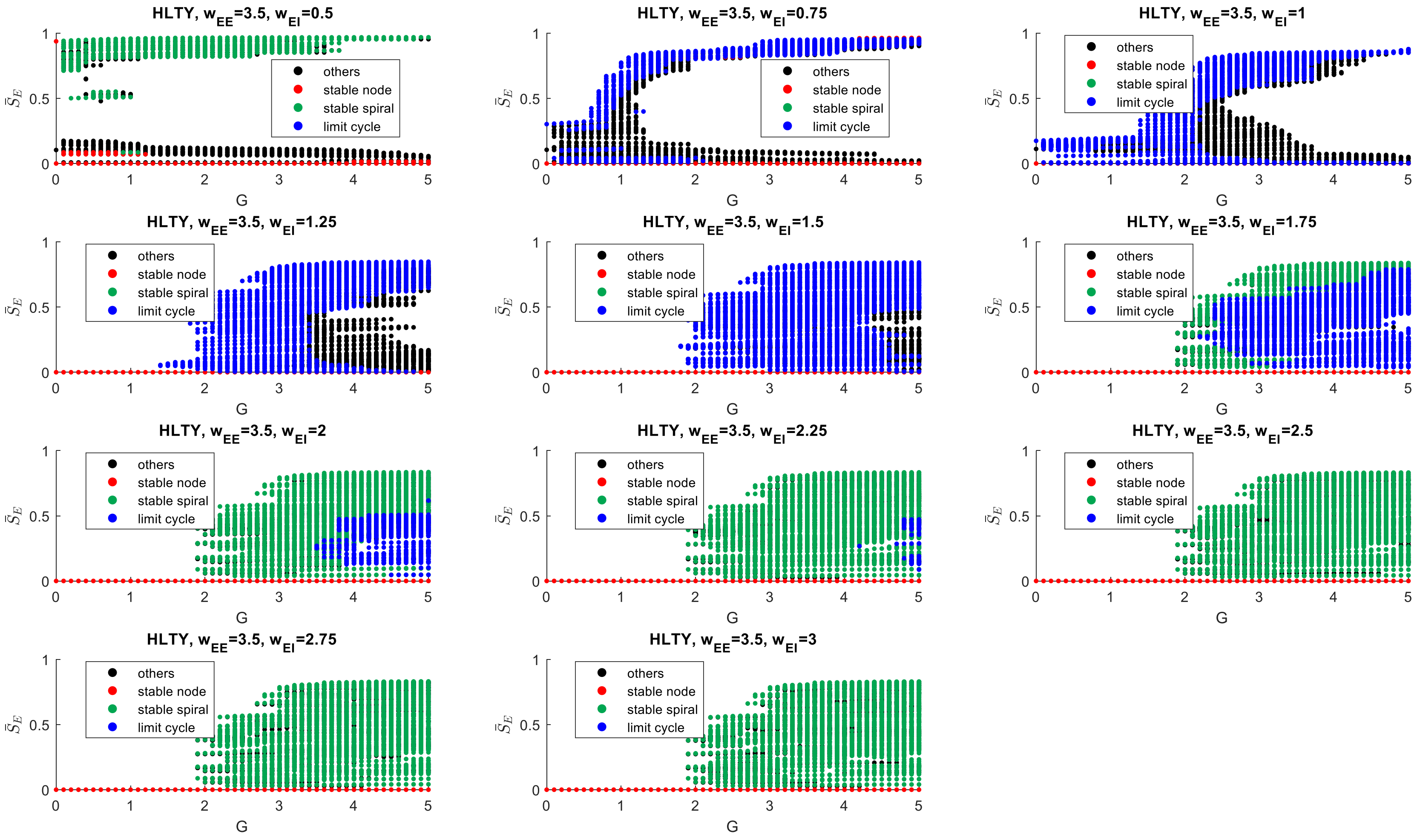


**Supplemental Fig. 21:** Population-level bifurcation plots for healthy controls ($W_{EE}$ = 3.5)

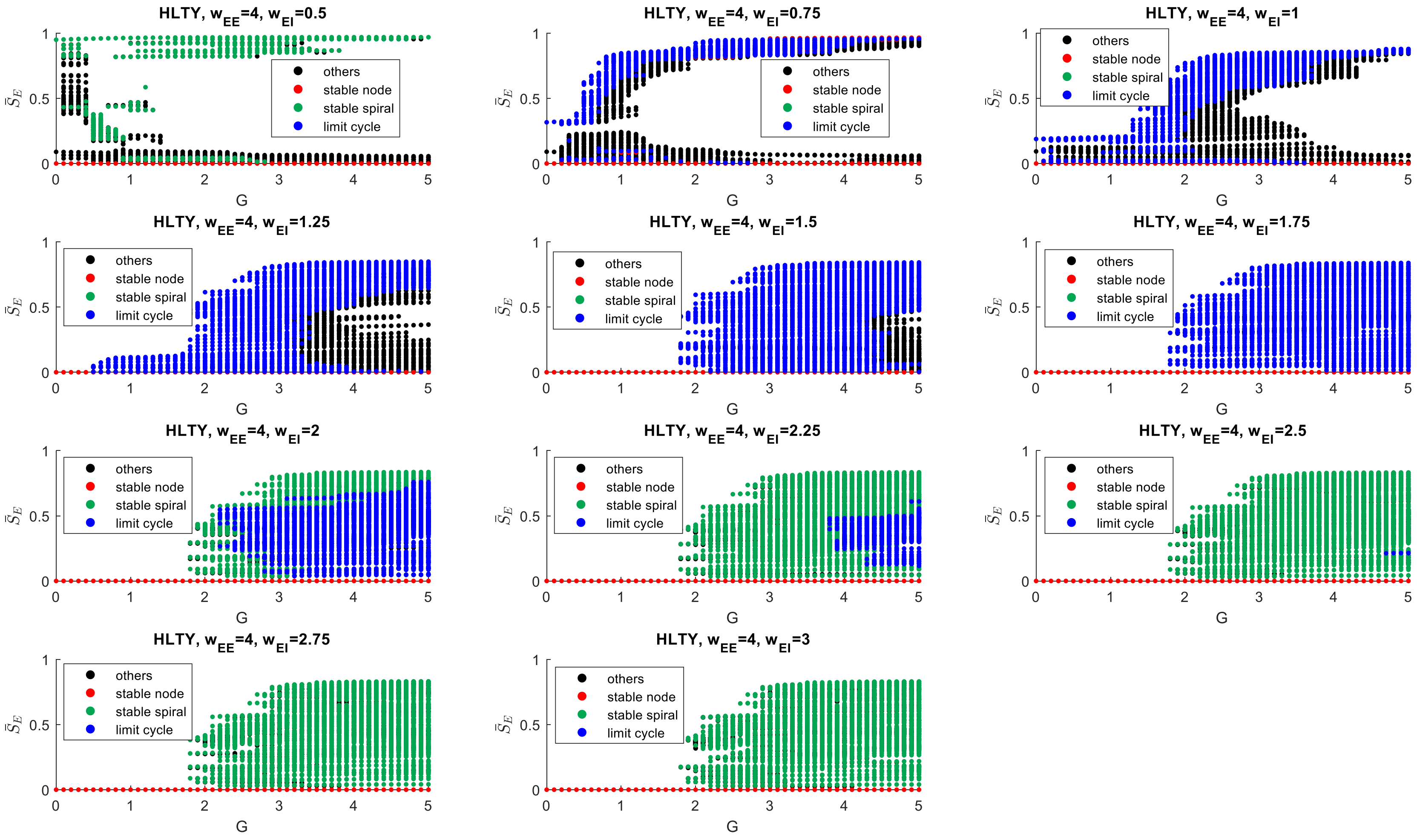


**Supplemental Fig. 22:** Population-level bifurcation plots for healthy controls ($W_{EE}$ = 4.0)

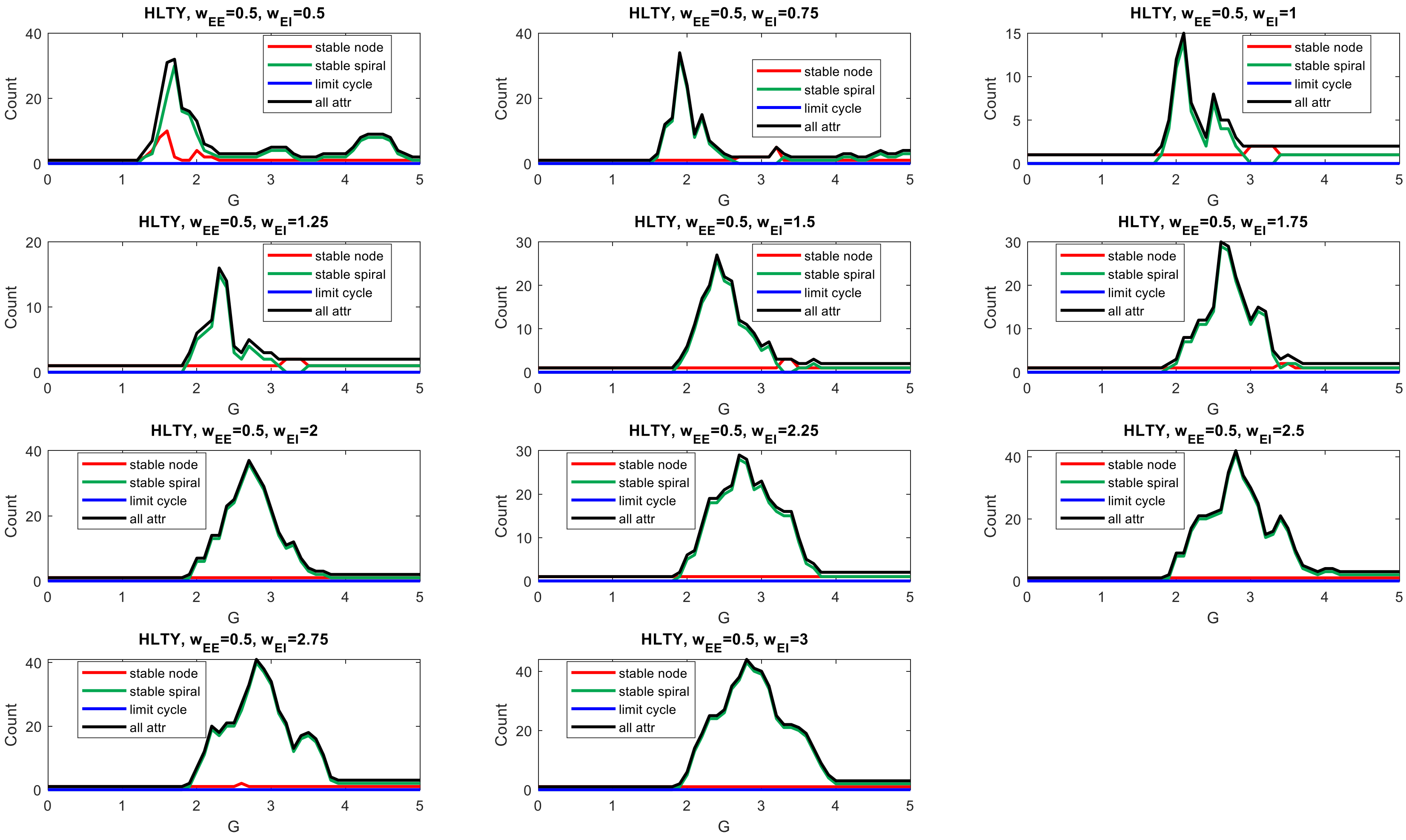


**Supplemental Fig. 23:** Change in attractor profile with global coupling (G) for healthy controls ($W_{EE}$ = 0.5)

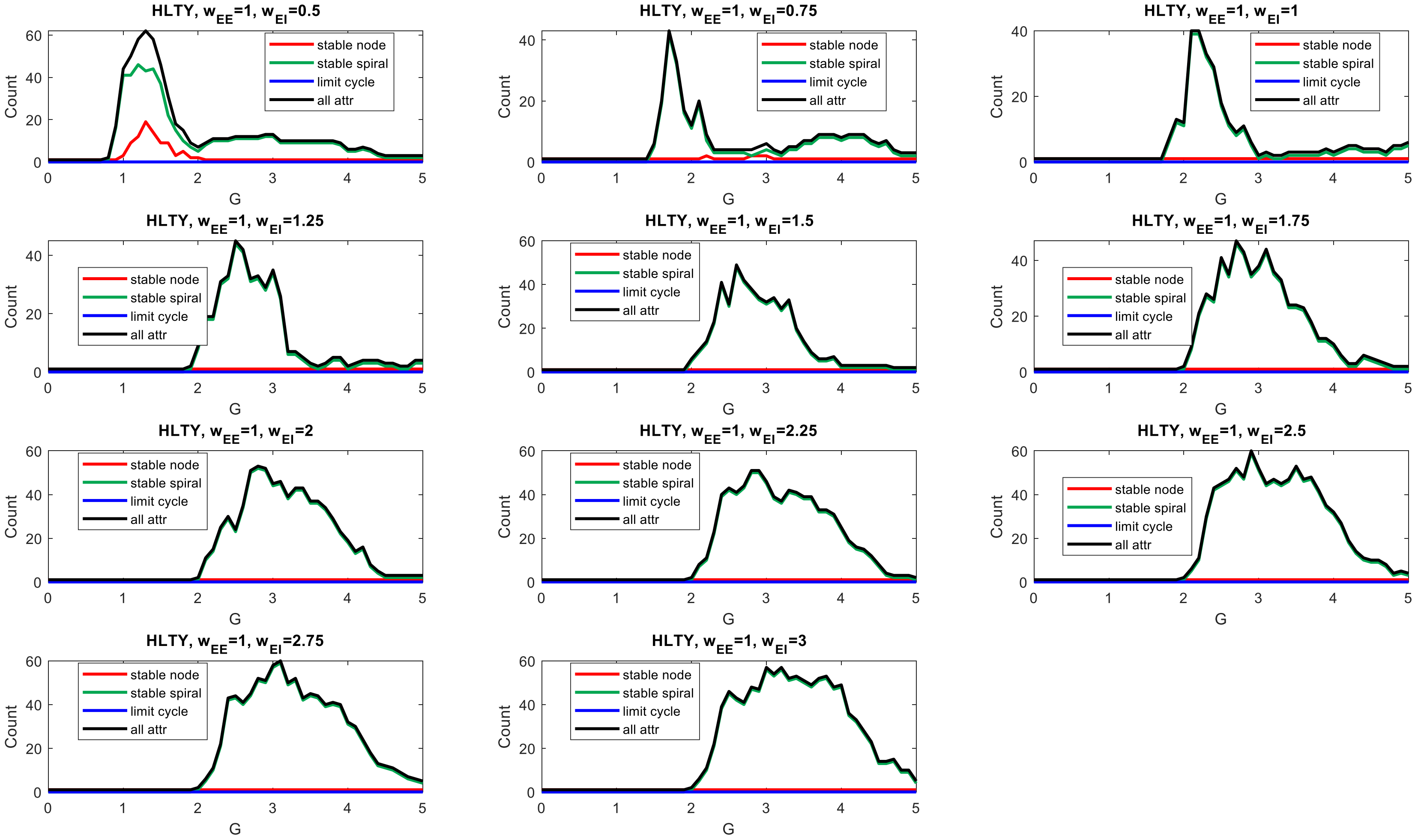


**Supplemental Fig. 24:** Change in attractor profile with global coupling (G) for healthy controls ($W_{EE}$ = 1.0)

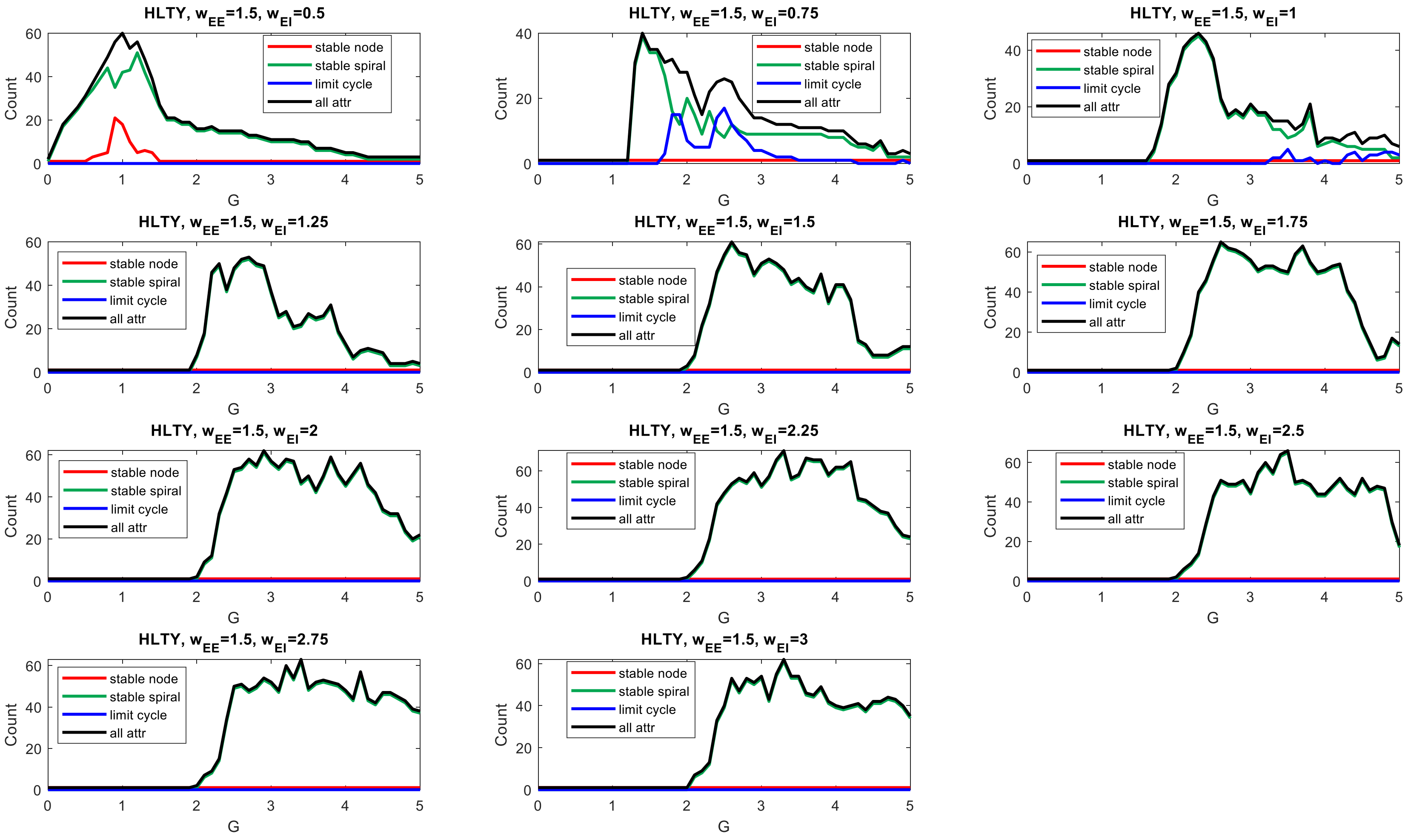


**Supplemental Fig. 25:** Change in attractor profile with global coupling (G) for healthy controls ($W_{EE}$ = 1.5)

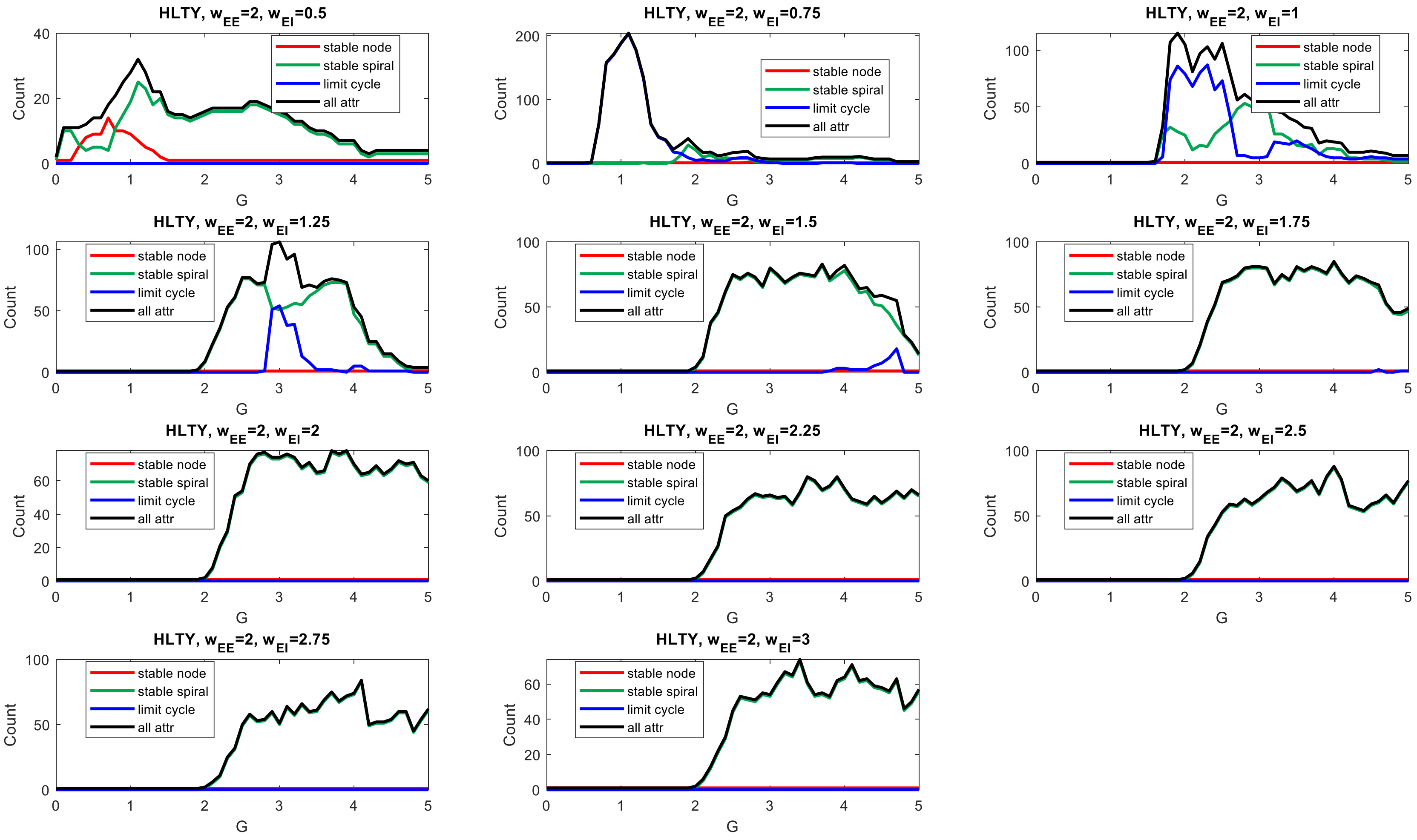


**Supplemental Fig. 26:** Change in attractor profile with global coupling (G) for healthy controls ($W_{EE}$ = 2.0)

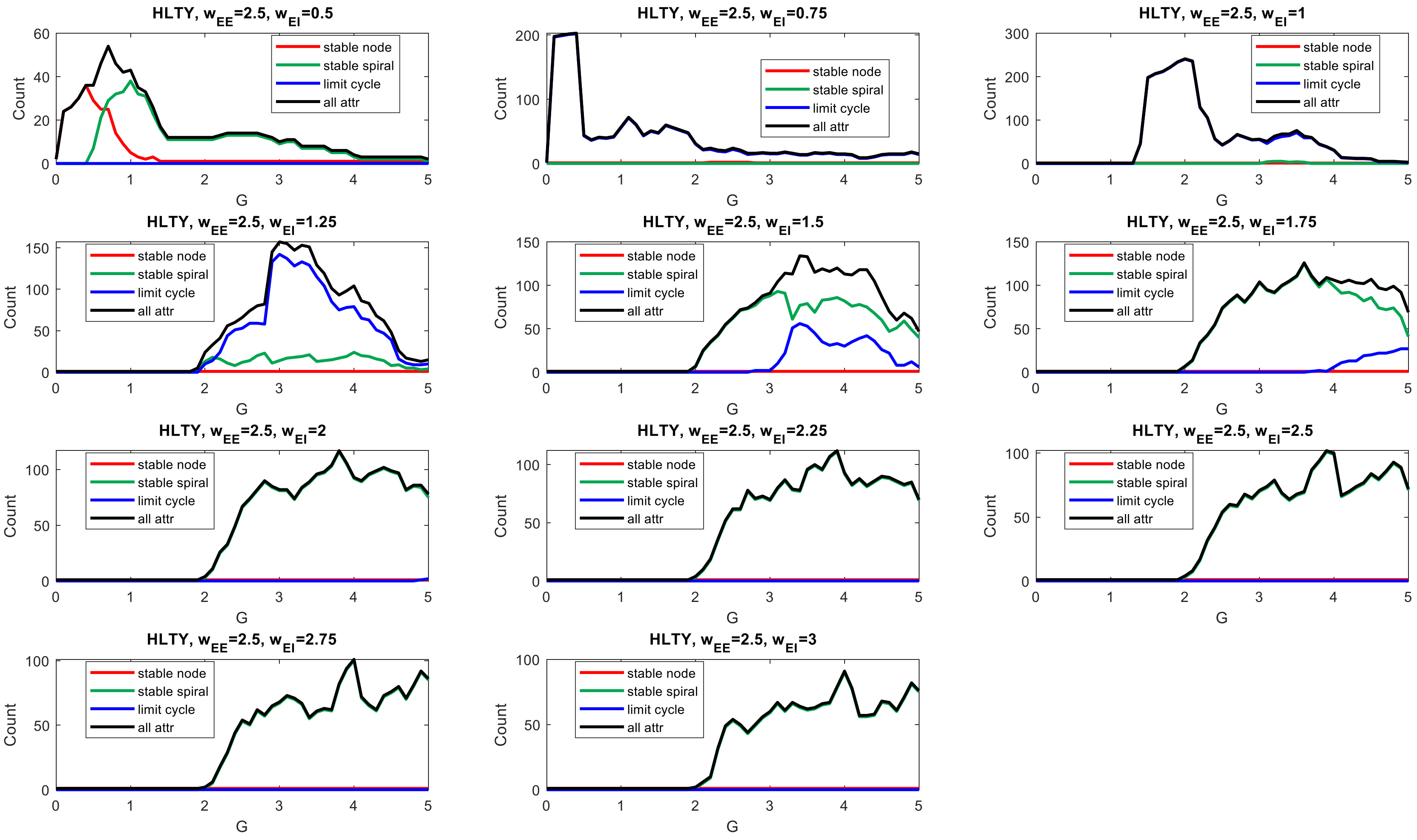


**Supplemental Fig. 27:** Change in attractor profile with global coupling (G) for healthy controls ($W_{EE}$ = 2.5)

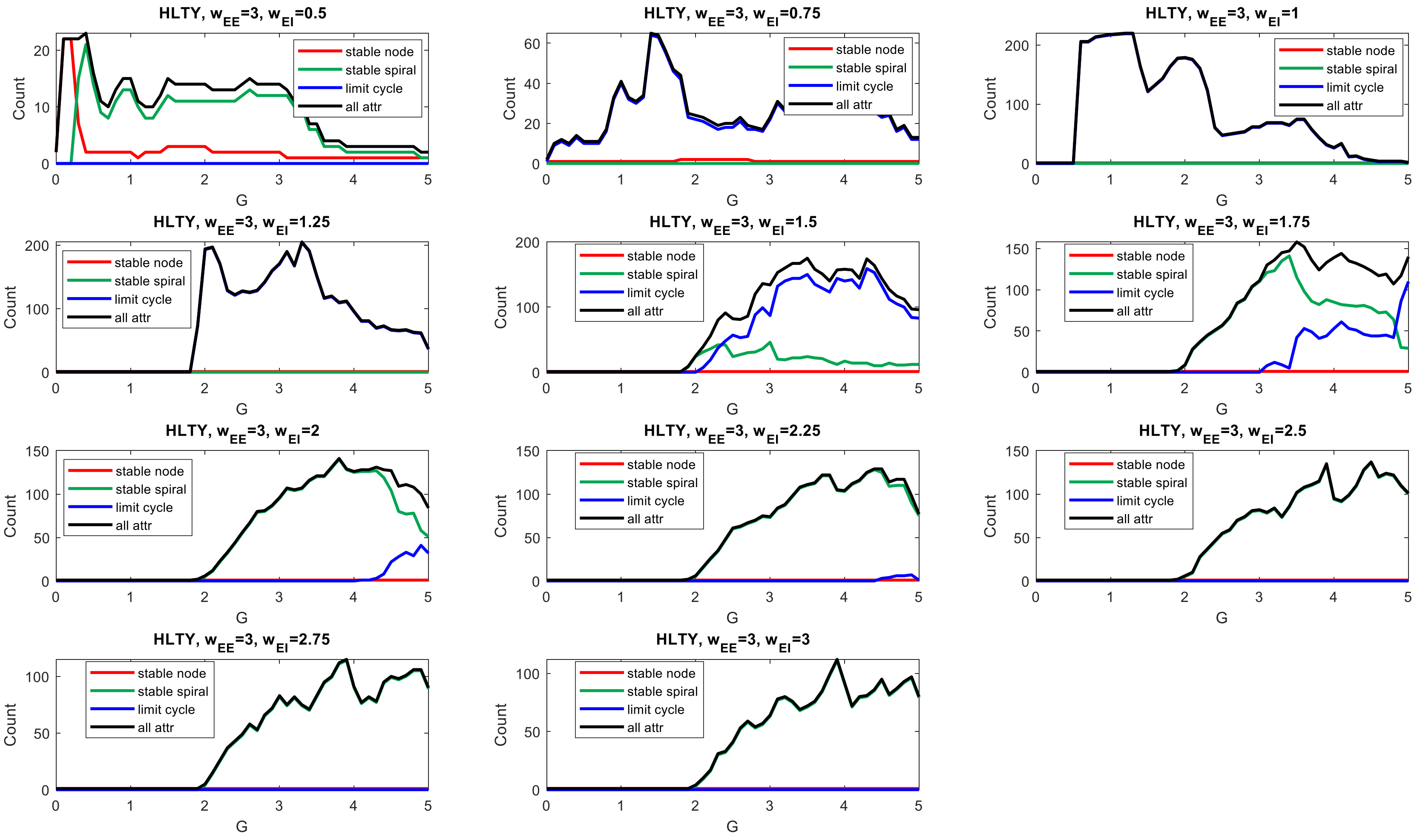


**Supplemental Fig. 28:** Change in attractor profile with global coupling (G) for healthy controls ($W_{EE}$ = 3.0)

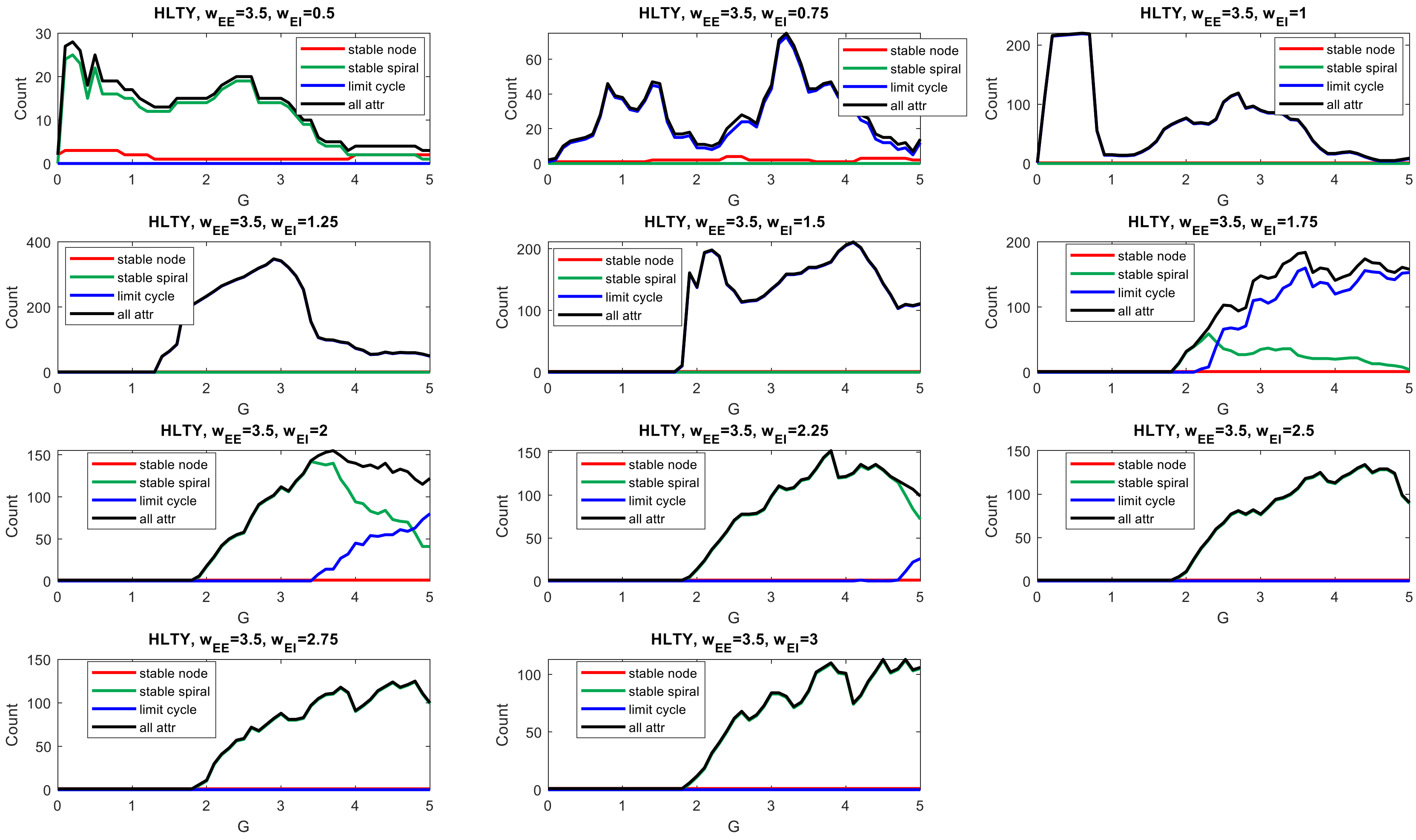


**Supplemental Fig. 29:** Change in attractor profile with global coupling (G) for healthy controls ($W_{EE}$ = 3.5)

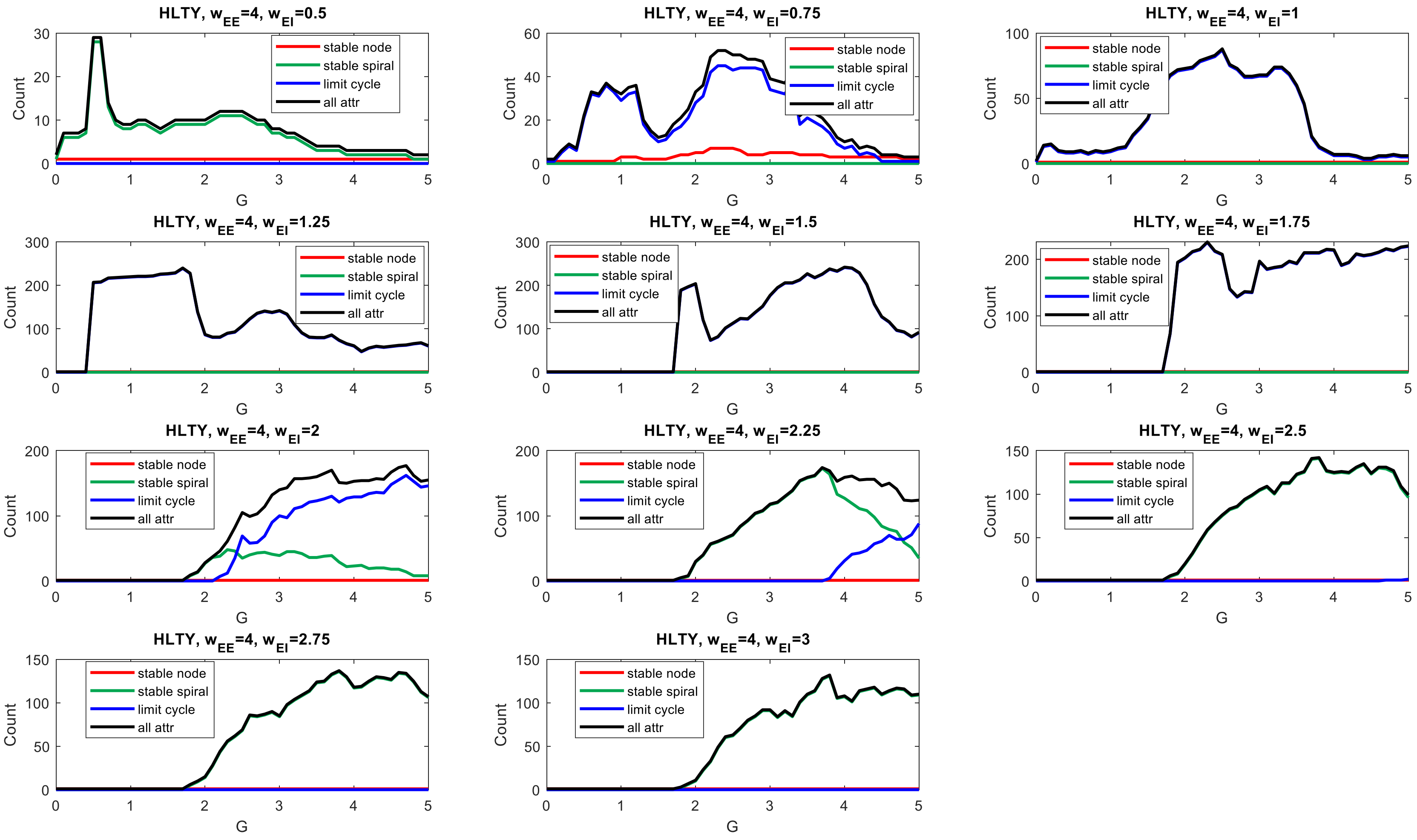


**Supplemental Fig. 30:** Change in attractor profile with global coupling (G) for healthy controls ($W_{EE}$ = 4.0)
